# Presynaptic inhibition switches hippocampal mossy fiber computations and limits vesicle pool plasticity

**DOI:** 10.64898/2026.11.23.768383

**Authors:** Peipeng Lin, Yuji Okamoto, Mojtaba R. Tavakoli, Johann G. Danzl, Peter Jonas, Katharina Lichter

## Abstract

Hippocampal mossy fiber synapses act as powerful teacher synapses in CA3 memory networks, yet how their detonation properties are regulated remains unclear. Combining presynaptic and paired recordings, electron and expansion microscopy, and optogenetic stimulation in mouse hippocampal slices, we show that both exogenously applied baclofen and ɣ-aminobutyric acid (GABA) synaptically released from inhibitory interneurons control transmission and plasticity at hippocampal mossy fiber synapses via presynaptic GABA_B_ receptors. Presynaptic inhibition markedly changes the computational properties of the synapse, switching it from a detonation into a subdetonation mode. Powerful modulation is enabled by a combination of effects on release probability, mediated by the suppression of presynaptic Ca^2+^ inflow, and on the readily releasable vesicle pool, associated with a reduced number and increased dispersion of docked synaptic vesicles at the ultrastructural level. Synaptically released GABA reverses established post-tetanic potentiation, suggesting interference with vesicle storage mechanisms potentially underlying short-term memory (“vesicle pool engram”). Our results reveal a previously unrecognized inhibitory-excitatory interaction at a central excitatory synapse, in which presynaptic inhibition both sets the computational properties of transmission and limits the persistence of vesicle pool engrams. (181 words)

**Teaser:** GABAergic control at mossy fiber synapses tunes detonation and limits vesicle pool plasticity in hippocampal memory circuits

## INTRODUCTION

Memory formation and retrieval rely on the hippocampal trisynaptic circuit, which relays information across three successive synapses, from the entorhinal cortex to the dentate gyrus, CA3, and CA1. The hippocampal mossy fiber synapse, connecting the granule cell layer to the CA3 subfield, is a key connection in this circuit (*1*). Several structural and functional properties suggest that this synapse is uniquely powerful, acting as a detonator that can fire a CA3 pyramidal neuron (PN) from the input of a single granule cell. At the structural level, the large size of the presynaptic terminals, the high density of synaptic vesicles, and the presence of multiple active zones (AZs) are thought to establish an efficient transmitter release mechanism (*2*, *3*). At the functional level, several short-term and long-term forms of presynaptic plasticity coexist at this synapse. These include facilitation, post-tetanic potentiation (PTP (*4*), and long-term potentiation (LTP (*5*)), which collectively increase synaptic efficacy by more than an order of magnitude. These unique structural and functional properties suggest that the mossy fiber synapse acts as teacher synapse, which drives memory storage in the recurrent collateral synapses of the downstream CA3 network (*1*, *6*). At the same time, however, they raise the question of how detonation is regulated at the level of AZs as sites of transmitter release and determinants of presynaptic strength (*7*, *8*).

Throughout the mammalian nervous system, excitation is controlled by postsynaptic inhibition, where interneurons release the transmitter GABA and activate GABA_A_ and GABA_B_ receptors (*9*, *10*). Alternatively, GABA released from inhibitory interneurons may act presynaptically (*11*). Presynaptic inhibition has been originally described in the spinal cord, where inhibitory transmitters released from interneurons suppress the release of excitatory transmitters from afferent fibers (*12*, *13*). In cortical circuits, however, the abundance, mechanisms, and significance of presynaptic inhibition are less clear. For mossy fiber synapses in the hippocampus, previous work suggested that presynaptic GABA_A_ receptors can regulate action potential (AP) propagation and transmitter release (*14*, *15*). In contrast, field potential recordings in the CA3 *stratum lucidum* indicated that GABA_B_ receptor activation markedly suppresses synaptic transmission at these synapses (*16*, *17*), pointing to a powerful modulatory control. Presynaptic GABA_B_ receptors, as G-protein coupled receptors, are generally thought to reduce release probability by inhibition of presynaptic Ca^2+^ channels (*18–20*), although effects on vesicle exocytosis have also been reported (*21*). Which of these mechanisms dominates at hippocampal mossy fiber terminals remains unresolved.

Presynaptic plasticity is widely thought to be mediated by changes in the probability of transmitter release (*5*). However, recent work suggests that many forms of presynaptic plasticity are generated by an increase in the size of the readily releasable vesicle pool (RRP) (*22*, *23*, *4*, *24*). In hippocampal mossy fiber synapses, where presynaptic plasticity is particularly prominent (*5*), both posttetanic potentiation (PTP) induced by high-frequency stimulation (HFS) and chemical potentiation by activation of the 3’,5’-cyclic adenosine monophosphate (cAMP) pathway are associated with an increase in RRP size (*4*, *25*). In both cases, functional changes are accompanied by an increased number of docked vesicles in AZs (*4*, *25*, *26*). Activation of GABA_B_ receptors is thought to inhibit cAMP production via G_i_ proteins (*27*). This suggests the possibility that activation of presynaptic GABA_B_ receptors inhibits presynaptic plasticity and specifically interferes with the induced structural changes in the vesicle pool (*4*). However, this hypothesis has not been directly tested.

It is widely thought that information in the brain is stored in the form of engrams, classically defined as the ‘physical and/or chemical changes that were elicited by learning and underlie the newly formed memory associations.’ (*28*, *29*). In principle, engrams may comprise changes at the molecular, cellular, and circuit level. Recent work demonstrated that changes in the organization of active zones and synaptic vesicle pools represent a substantial part of the engram, contributing to formation and retention of memories (*30*, *31*). How inhibition shapes engrams remains unclear. It has been suggested that GABAergic neurons generate independent inhibitory engrams (*32*, *33*). Alternatively, it is possible that inhibition affects excitatory engrams, for example by regulating the size of the releasable vesicle pool. Direct experimental approaches are needed to distinguish between these possibilities.

To address these questions, we examined the biophysical and structural mechanisms of presynaptic inhibition at hippocampal mossy fiber bouton (MFB)–CA3 PN synapses. The giant presynaptic terminals enabled us to resolve presynaptic structure-function relationships at single-vesicle resolution by combining subcellular recordings (*34*) with functional two-dimensional (2D) and three-dimensional (3D) electron microscopy of near-to-natively cryo-immobilized and freeze-substituted hippocampal MFBs (“pharm-and-freeze”, “flash-and-freeze”) (*26*, *35*). Our results suggest that presynaptic inhibition shifts the mossy fiber synapse from a detonation into a sub-detonation mode, demonstrating that presynaptic inhibition switches the synapse from a logic “OR gate” into an “AND gate” (*36*, *37*). Furthermore, our data indicate that activation of presynaptic GABA_B_ receptors not only suppresses release probability by inhibiting presynaptic Ca^2+^ channels, but also reduces the size of the RRP, both when assessed functionally and structurally, thereby halting short-term plasticity and limiting memory storage (*4*). Finally, using super-resolution expansion microscopy, we identified a close spatial relationship between GABAergic terminals and mossy fiber synapses in CA3 *stratum lucidum*. This provides a structural basis for how synaptically released GABA can modulate transmitter release and short-term plasticity.

## RESULTS

### GABA_B_ receptor activation suppresses MFB–CA3 PN synaptic transmission

To dissect the biophysical mechanisms of presynaptic inhibition in hippocampal MFB–CA3 synapses, we performed paired recordings between single mossy fiber terminals and postsynaptic CA3 pyramidal cells in mice (Fig. 1). MFBs were stimulated in the cell-attached configuration to avoid cytoplasmic washout, and postsynaptic responses were recorded in the whole-cell configuration (Fig. 1A). Bath application of 20 µM of the GABA_B_ receptor agonist R-(−)-baclofen reduced the amplitude of the first excitatory postsynaptic current (EPSC_1_) in a 50-Hz train of 15 stimuli to 10.14 ± 2.06% of the control value (Ctrl: 358.3 ± 80.6 pA; baclofen: 27.1 ± 4.9 pA; Fig. 1, B to D; 11 MFB−CA3 PN paired recordings; *p* = 0.001). While baclofen application increased the proportion of failures and the EPSC_1_ latency, it reduced the EPSC_1_ 20–80% rise time and decay time constant, consistent with a presynaptic effect (Fig. 1E to H). To address whether presynaptic GABA_B_ receptors were tonically activated, we further tested the effects of 5 µM of the GABA_B_ receptor antagonist CGP 55845 (Fig. S1; Table S1). In contrast to the marked effects of the agonist, the antagonist had no significant effects, arguing against tonic receptor activation. Furthermore, CGP 55845 did not significantly enhance EPSCs during HFS, as would be expected if activation of a single MFB recruited interneurons and induced disynaptic inhibition (Fig. S1; Table S1). Taken together, these results suggest that presynaptic GABA_B_ receptors control transmission at MFB–CA3 synapses, but do not seem to be activated under resting conditions or via disynaptic inhibition.

**Fig. 1.**
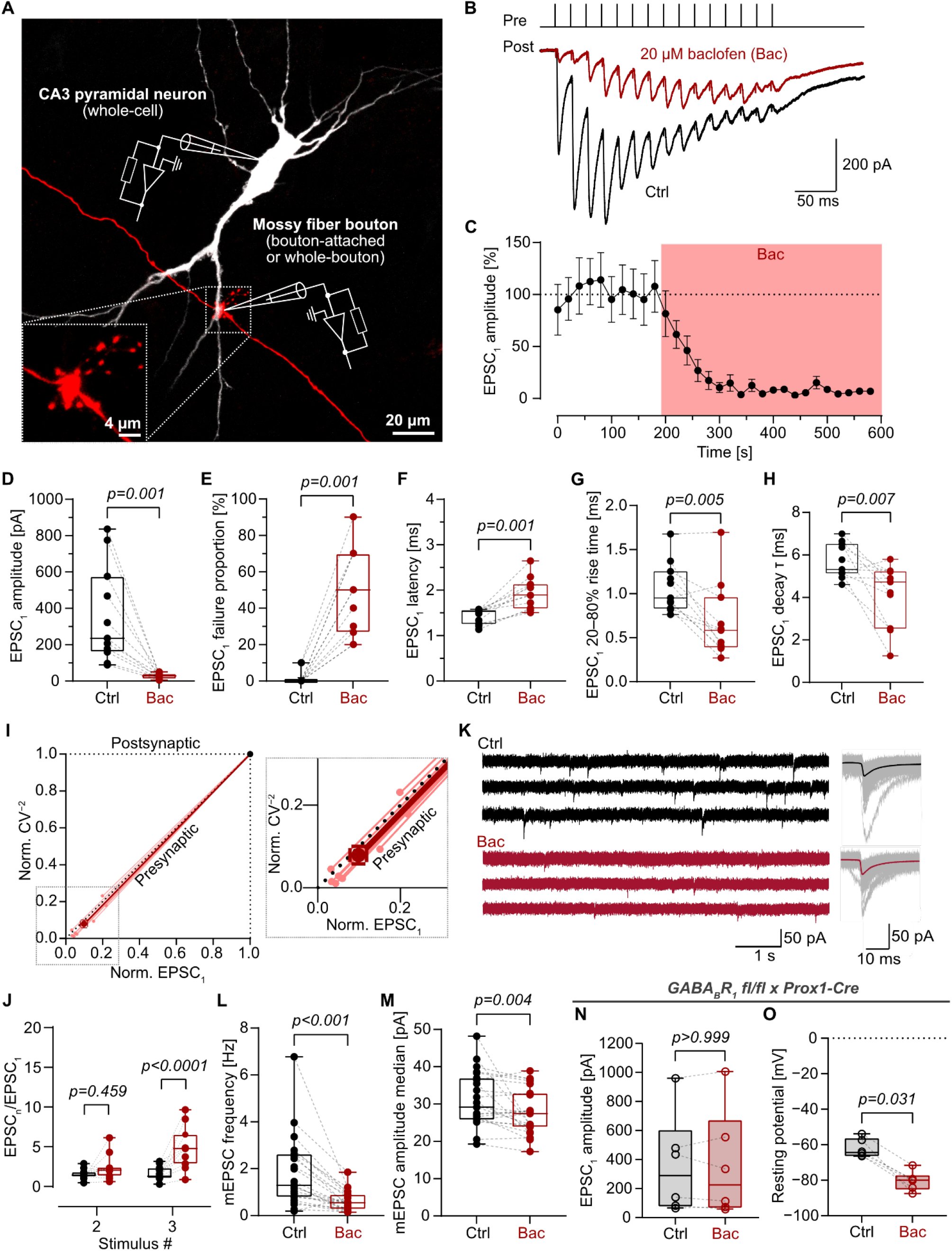
The selective GABA_B_ receptor agonist baclofen suppresses MFB–CA3 PN synaptic transmission via presynaptic GABA_B_ receptors. **(A)** Maximum intensity Z-projection of a confocal micrograph of paired pre-postsynaptic recording from a biocytin-labeled mossy fiber bouton (MFB, red) and its synaptically connected CA3 pyramidal neuron (PN, white). **(B)** Representative traces of EPSCs evoked by a 50-Hz train of 15 stimuli in standard ACSF (Ctrl, black) and in the presence of 20 µM GABA_B_ receptor agonist baclofen (Bac, red) from the same MFB–CA3 PN pair. Traces were averaged from 10 sweeps in each condition. **(C)** Normalized EPSC_1_ peak amplitudes (to the average EPSC_1_ peak amplitude from Ctrl) plotted against experimental time. Bac application, light red shading. **(D–H)** Summary box plots of EPSC_1_ peak amplitude, EPSC_1_ failure proportion, EPSC_1_ latency, EPSC_1_ 20–80% rise time, and EPSC_1_ decay time constant τ. Data points from the same MFB–CA3 PN pairs are connected by dashed lines. **(I)** Normalized coefficient-of-variation raised to power of −2 (CV^−2^) plotted against normalized change in EPSC_1_ peak amplitude (left) with zoomed inset on data points from individual MFB–CA3 PN pairs (pink data points, right). **(J)** Summary box plot of EPSC_n_/EPSC_1_ for stimulus number 2 and 3 from the stimulus train in Ctrl and Bac. Data in C–J from N = 11 MFB–CA3 PN pairs. **(K)** Representative traces of mEPSC recordings in Ctrl and Bac in CA3 PNs (left). Individual mEPSC events are detected and aligned at an expanded time scale (gray), overlaid with the average mEPSC in Ctrl and baclofen (right). **(L, M)** Summary box plot of mEPSC frequency and amplitude median in Ctrl and Bac. N = 23 CA3 PNs. **(N, O)** Summary box plots of EPSC_1_ peak amplitude, and resting membrane potential of CA3 PNs in Ctrl and Bac, measured in GABA_B_1 fl/fl x Prox1-Cre mice. N = 6 MFB–CA3 PN pairs.

Next, we asked whether the effects of GABA_B_ receptor activation on synaptic transmission at individual mossy fiber synapses are mediated pre- or postsynaptically. First, we performed a coefficient-of-variation analysis of the paired recordings obtained under baclofen application in wild-type animals, plotting the normalized coefficient-of-variation raised to −2 power (CV^−2^) against the normalized EPSC_1_ amplitude (*38*). The data points from 11 paired recordings were located on the identity line, indicating a presynaptic site of action (Fig. 1I; normalized EPSC_1_: 0.10 ± 0.02; normalized CV^−2^: 0.08 ± 0.02). Second, we examined the effects of baclofen on the synaptic dynamics. Although baclofen did not significantly affect the paired pulse ratio (EPSC_2_/EPSC_1_), EPSC_3_/EPSC_1_ was significantly enlarged (Fig. 1J; Table S1). Third, we tested for the effect of baclofen on miniature EPSCs (mEPSCs) in CA3 PNs and found a reduced frequency, and a small but significant reduction in amplitude, corroborating a presynaptic locus for baclofen-mediated presynaptic inhibition (Fig. 1, K to M; frequency – Ctrl: 1.84 ± 0.31 Hz; baclofen: 0.63 ± 0.09 Hz; *p* < 0.001; amplitude (median) – Ctrl: 31.3 ± 1.5 pA; baclofen: 27.8 ± 1.3 pA; *p* = 0.004; 23 CA3 PN recordings), in line with previous findings (*39*). To estimate the effects of baclofen on mEPSC events of putative mossy fiber origin, we selectively analyzed the frequency and amplitude of mEPSCs with fast kinetics (20–80% rise time < 1 ms), as previously described (*39*), and obtained similar results (Fig. S2; Table S2).

Finally, to unequivocally determine the locus of GABA_B_ receptor action on mossy fiber transmission, we selectively deleted the relevant GABA_B_ receptor 1 subunit (*16*) from dentate gyrus granule cells by generating a conditional knockout (cKO) mouse (GABA_B_R_1_ fl/fl x Prox1-Cre (*26*, *40*)). In cKO animals, baclofen no longer affected EPSC_1_ amplitudes (Fig. 1N, 6 MFB−CA3 PN paired recordings). The structure of the mossy fiber tract including MFBs was unaffected in the cKO (Figure S3), and postsynaptic GABA_B_ receptor function was also preserved, with baclofen-induced membrane hyperpolarization and accompanying positive shift in holding current (∼100–200 pA) comparable to wild-type animals (Fig. 1O; Fig. S3; Table S1). Taken together, GABA_B_ receptor activation markedly suppressed mossy fiber transmission via presynaptic mechanisms.

### Presynaptic GABA_B_ receptor activation changes single-synapse computations from detonation to subdetonation

A hallmark property of mossy fiber transmission is the efficacy of the synapse, which generates reliable spiking in postsynaptic CA3 neurons following conditional or plasticity-dependent detonation (*1*, *41*, *42*). To investigate how presynaptic GABA_B_ receptor activation affects synaptic computation between dentate gyrus and CA3, we examined the detonation properties of the synapse by measuring effects of baclofen on excitatory postsynaptic potentials (EPSPs, Fig. 2). Single presynaptic MFBs were stimulated by 50-Hz trains of three stimuli, and postsynaptic responses of CA3 PNs were recorded as EPSPs under current-clamp conditions (Fig. 2A). To avoid any confounding effects of the baclofen-induced hyperpolarization of the postsynaptic neuron through activated G protein-coupled inwardly rectifying potassium (GIRK) channels (Fig. 1O) (*43*, *44*), the postsynaptic membrane potential was held close to the average resting potential of CA3 PNs by constant current injection (*45*).

**Fig. 2.**
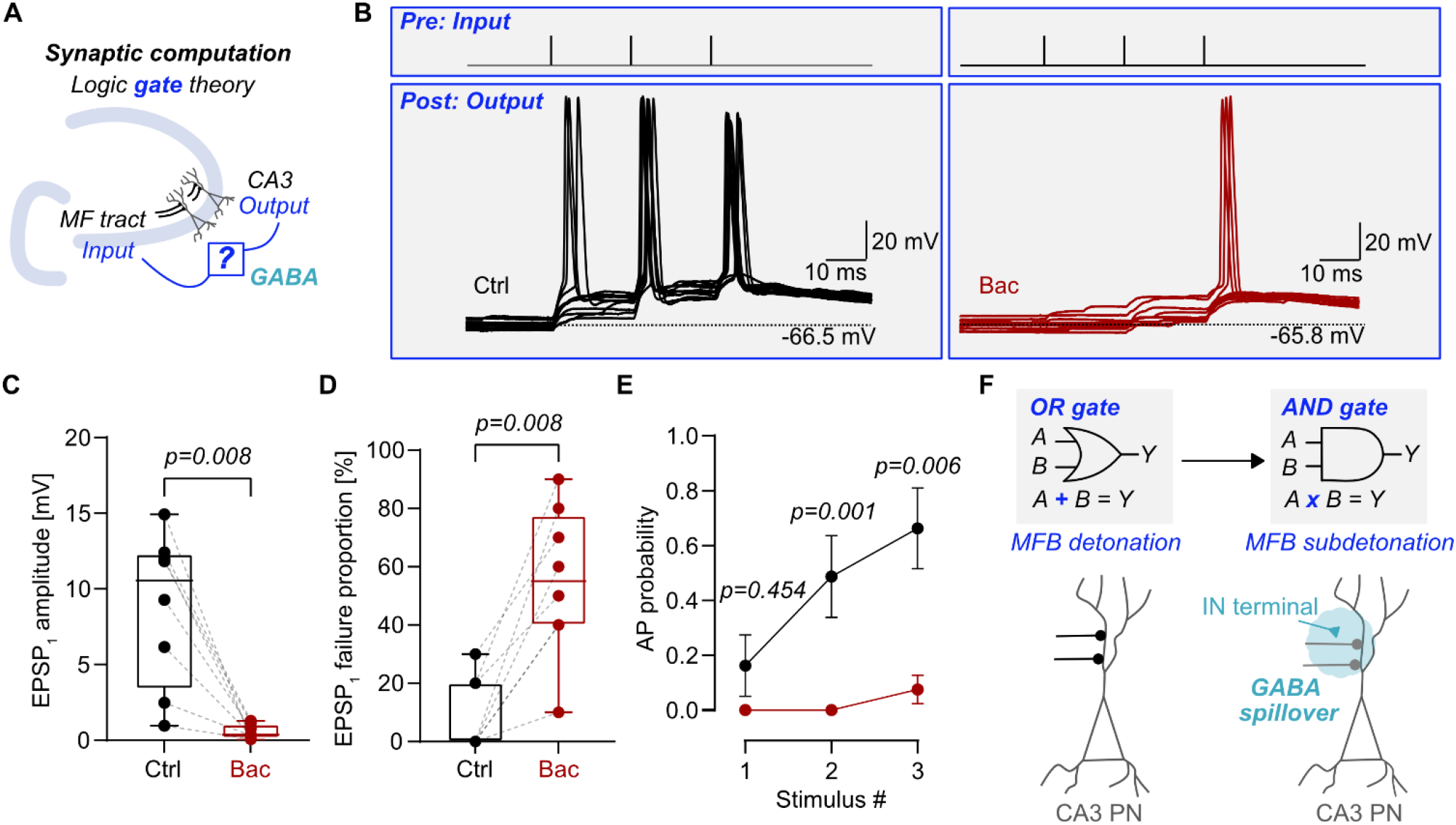
Presynaptic GABA_B_ receptor activation switches MFB–CA3 PN synapses from detonation into subdetonation mode. **(A)** Schematic of synaptic computation between individual MFBs (input) and CA3 PNs (output), framed as logic gates, illustrating how spillover GABA and presynaptic inhibition interferes with CA3 network properties. **(B)** Representative traces of presynaptic stimuli (input) and postsynaptic EPSPs and APs (output) evoked by 50-Hz trains of 3 stimuli in Ctrl and Bac. **(C, D)** Summary box plots of EPSP_1_ peak amplitude **(C)** and EPSP_1_ failure proportion **(D)** in Ctrl and Bac. **(E)** Probability of AP firing at each stimulus in Ctrl and Bac. Data in B–E from N = 8 MFB–CA3 PN pairs. **(F)** Presynaptic inhibition reconfigures synaptic computations at CA3 PNs, switching from an “OR gate” (left) to an “AND gate” (right). “OR gate”: A single MFB detonation (A or B) is sufficient to evoke an AP (Y) in a CA3 PN. “AND gate”: Spillover GABA from a proximal interneuron terminal switches MFBs from detonation to subdetonation mode, requiring coincident activation of both MFBs (A and B) to evoke a postsynaptic AP (Y).

Under control conditions, trains of stimuli reliably evoked APs in the postsynaptic CA3 PN (Fig. 2B). In contrast, after application of baclofen, AP probability was markedly reduced. The amplitude of EPSP_1_ in a 50-Hz train of three stimuli was changed from 8.75 ± 1.79 mV (Ctrl) to 0.54 ± 0.17 mV (baclofen) (Fig. 2C; Table S3; *p* = 0.008; 8 MFB−CA3 PN paired recordings). These changes were associated with a significantly increased proportion of failures (Fig. 2D; *p* = 0.008). In parallel, the probability of AP firing was markedly suppressed in baclofen condition (Fig. 2E; probability of first AP – Ctrl: 0.16 ± 0.11; baclofen: 0 ± 0; *p* = 0.454; probability of second AP – Ctrl: 0.49 ± 0.15; baclofen: 0 ± 0; *p* = 0.001; probability of third AP – Ctrl: 0.66 ± 0.15; baclofen: 0.08 ± 0.05; *p* = 0.006). Thus, GABA_B_ receptor activation markedly changed single-synapse computations, converting the mossy fiber synapse from detonation to subdetonation (Fig. 2F) (*41*, *42*). Functionally, this shifts the synapse from an “OR gate”, in which a single MFB can induce postsynaptic activity, to an “AND gate”, in which firing requires several MFBs to be active together (Fig. 2F) (*42*).

### Presynaptic GABA_B_ receptor activation reduces the size of the releasable vesicle pool in single mossy fiber terminals

Previous work using field potential recordings suggested that a presynaptic GABA_B_ receptor activation reduces release probability at mossy fiber synapses (*16*). However, the effects of this modulation on synaptic efficiency may be more complex. We therefore analyzed the biophysical mechanisms underlying presynaptic inhibition by GABA_B_ receptors at the single bouton level (Fig. 3). To quantify vesicular release probability (P_v_) and RRP size, we performed cumulative EPSC analysis (*4*, *46*). MFBs were repetitively stimulated in the cell-attached configuration, and EPSCs in postsynaptic CA3 PNs were recorded. EPSC peak amplitudes were cumulatively plotted against time, and the last four responses were fit by linear regression (Fig. 3A).

**Fig. 3.**
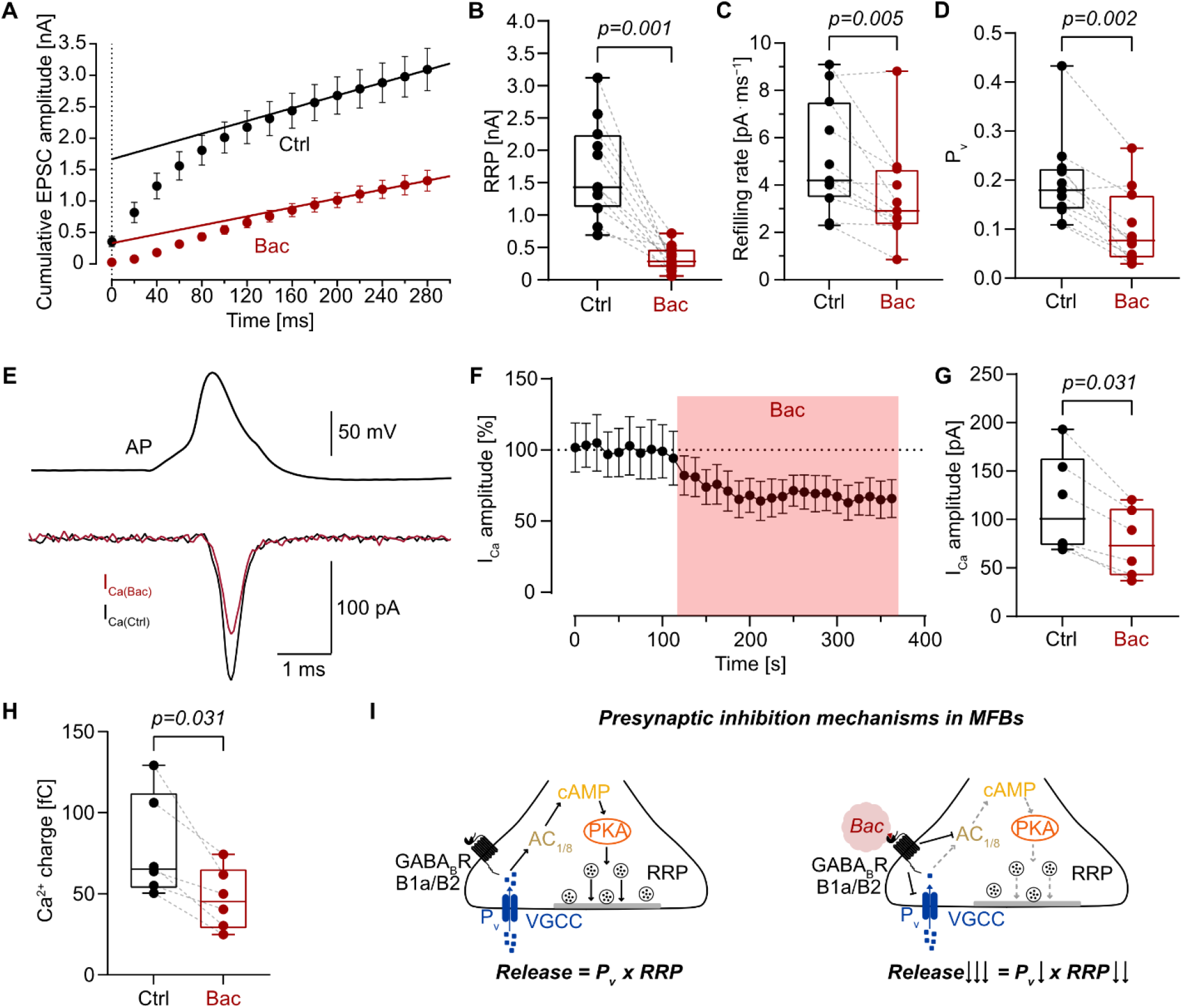
Baclofen substantially reduces RRP size and, to a small extent, vesicular release probability and presynaptic Ca^2+^ current. **(A)** Cumulative plot of EPSC peak amplitudes evoked by a 50-Hz train of 15 stimuli in Ctrl and Bac. RRP size, refilling rate, and P_v_ were estimated by linear regression of the last four data points and back-extrapolation to time 0. **(B–D)** Summary box plots of RRP size, refilling rate, and P_v_ in Ctrl and Bac. Data in A–D from N = 11 MFB–CA3 PN pairs. **(E)** Representative traces of presynaptic Ca^2+^ current evoked in an MFB by a voltage command of a previously recorded AP waveform in Ctrl and Bac. **(F)** Normalized presynaptic Ca^2+^ current peak amplitudes (to the average presynaptic Ca^2+^ current peak amplitude in Ctrl) plotted against experimental time. Bac application, light red shading. **(G, H)** Summary box plots of presynaptic Ca^2+^ current peak amplitude and Ca^2+^ total charge in Ctrl and Bac. Data in F–H from N = 6 MFBs. **(I)** Presynaptic mechanisms of evoked vesicular release under control (left) and during GABA_B_ receptor activation by baclofen (Bac; red cloud). GABA_B_ receptor activation diminishes RRP size and vesicle docking, and to small extent release probability and presynaptic Ca^2+^ current, possibly via the cAMP-PKA pathway. Together, presynaptic inhibition decreases evoked vesicular release at the MFB–CA3 PN synapse.

Comparison of recordings before and after application of baclofen revealed that the size of the RRP, estimated as the intercept of the fit line with the vertical axis, was substantially reduced in the presence of baclofen (Fig. 3B; Ctrl: 1.70 ± 0.23 nA; baclofen: 0.33 ± 0.06 nA; *p* = 0.001; 11 MFB−CA3 PN paired recordings). For pool analysis, only paired recordings which reached steady-state depression larger than 60% relative to the EPSC_1_ were included. Furthermore, the refilling rate, determined from the slope of the fit line, was diminished (Fig. 3C; Ctrl: 5.18 ± 0.72 pA ms^−1^; baclofen: 3.55 ± 0.63 pA ms^−1^; *p* = 0.005). In addition, the P_v_, measured as EPSC_1_ / RRP, was markedly reduced (Fig. 3D; Ctrl: 0.20 ± 0.03; baclofen: 0.10 ± 0.02; *p* = 0.002). Similar results were obtained when the refilling rate was assumed to be dependent on prior release of synaptic vesicles (Fig. S4; Table S1) (*47*). While both methods for estimating the RRP size reflect the decrement of the pool during stimulation rather than its absolute size (*48*), they complementarily indicate a reduction under baclofen application. In addition, RRP size estimates were unlikely to be confounded by saturation or desensitization of postsynaptic α-amino-3-hydroxy-5-methyl-4-isoxazolepropionic acid (AMPA) receptors during HFS, based on previous findings (*4*) and AMPA receptor subunit composition in CA3 PNs (*49*). Ultimately, baclofen application did not change RRP, refilling rate and P_v_ in cKO animals (Fig. S3; Table S1). Taken together, these results indicate that GABA_B_ receptor activation not only suppresses P_v_, but also substantially reduces the size of the RRP (Fig. 3I).

To further differentiate biophysical mechanisms underlying the effects on release probability, we analyzed the effects of GABA_B_ receptor activation on presynaptic Ca^2+^ currents (*50*) (Fig. 3, E to H). Hippocampal MFBs were recorded in the whole-cell voltage-clamp configuration, and presynaptic APs, previously recorded in MFBs under current-clamp conditions, were applied as voltage-clamp command waveforms (*51*). Baclofen reduced the peak amplitude of the AP-evoked Ca^2+^ current from 115.3 ± 20.9 pA (Ctrl) to 75.5 ± 14.2 pA (baclofen) (Fig. 3, E to G; *p* = 0.031; 6 MFB recordings). In addition, the Ca^2+^ charge was reduced from 78.53 ± 12.95 fC (Ctrl) to 46.97 ± 7.71 fC (baclofen), reflecting a reduced total Ca^2+^ influx (Fig. 3H, *p* = 0.031). In combination with the previously determined power coefficient of Ca^2+^ dependence of synaptic transmission at hippocampal mossy fiber synapses (1.77) (*52*), these results closely predicted the estimated changes in P_v_ (Table S5). Thus, GABA_B_ receptor-mediated suppression of presynaptic Ca^2+^ influx explains the reduction in release probability at hippocampal mossy fiber synapses (Fig. 3I).

### Presynaptic GABA_B_ receptor activation reduces the docked vesicle pool and selectively depletes large clear vesicles at individual mossy fiber AZs

The RRP on a functional level is often equated with the docked vesicle pool obtained by structural analysis (*53*, *54*). Consistent with this idea, activation of the cAMP pathway in MFBs increases both RRP and docked vesicle number in a correlated manner (*25*). Whether such a correlation also holds for inhibitory neuromodulation, such as GABA_B_ receptor-mediated presynaptic inhibition, remains unclear. To test this hypothesis, we compared the nanoscale organization of AZs, central dynamic structures involved in neurotransmitter release and fine-tuning of synaptic efficiency (*7*, *8*), in control conditions and in the presence of baclofen (Fig. 4; Fig. 5; Fig. S5; Fig. S6).

**Fig. 4.**
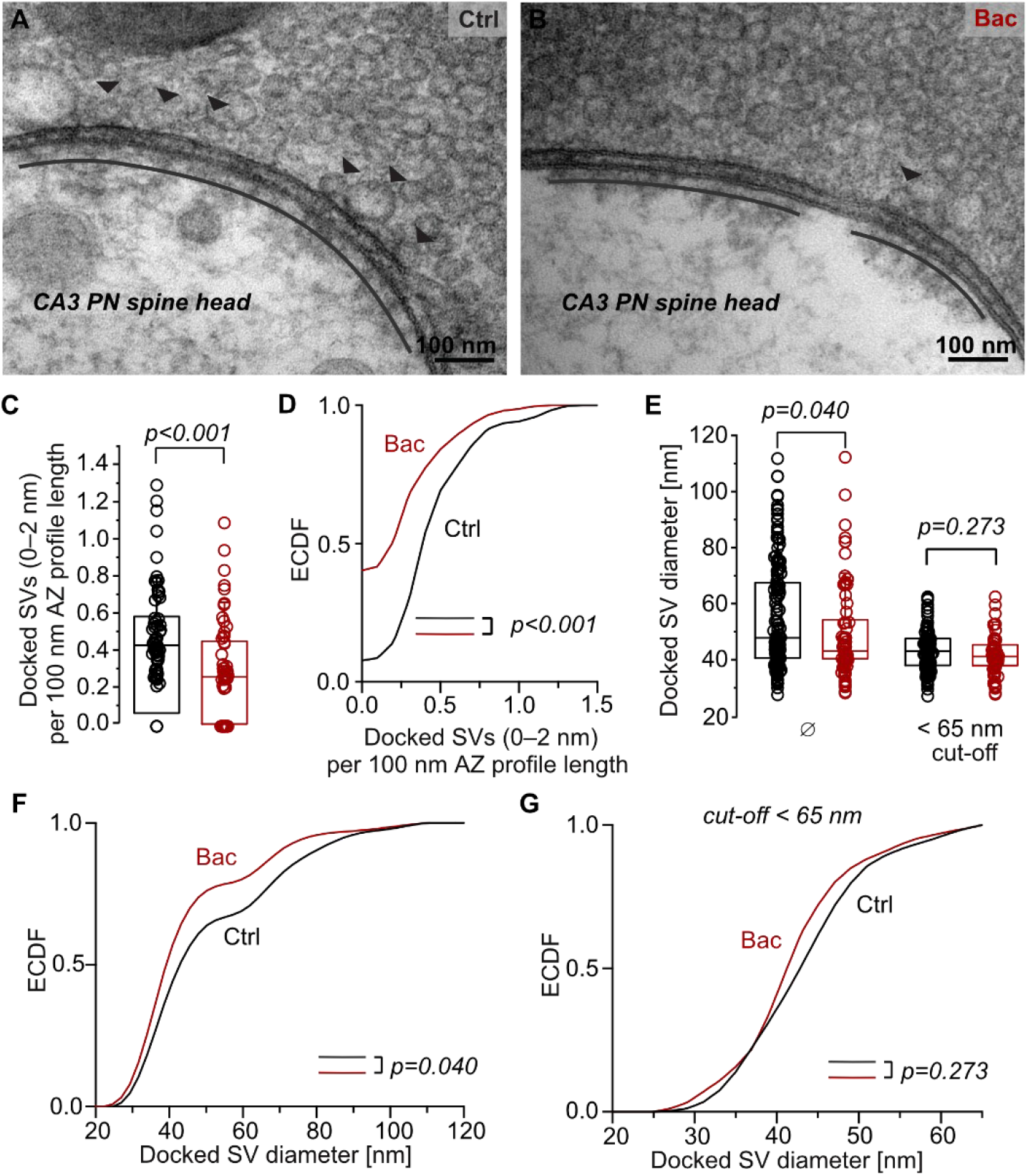
Baclofen reduces the number of docked vesicles and selectively depletes large clear vesicles. **(A, B)** Representative 2D electron micrographs of 100-nm ultrathin sections from high-pressure frozen acute brain slices in Ctrl **(A)** and Bac **(B,** ‘pharm-and-freeze’), highlighting presynaptic active zones (AZs), defined as structural compartments with docked synaptic vesicles (SVs, arrowheads) opposite to the postsynaptic density (extent indicated by black lines), of MFBs in *stratum lucidum* of the CA3b subregion. **(C, D)** Summary box plot **(C)** and empirical cumulative distribution function (ECDF) plot **(D)** of the number of docked (0–2 nm distance to the AZ membrane) per 100 nm AZ profile length in Ctrl and Bac. **(E–G)** Summary box plots and ECDF plots of the diameter of docked SVs (0–5 nm) in Ctrl and Bac, for all docked SVs and only including SVs with a diameter < 65 nm. Data in C–G from N_Ctrl_ = 65 AZs (2 mice) and N_Bac_ = 62 AZs (3 mice), data in E–G from N_Ctrl_ = 146 docked vesicles / 101 docked vesicles (< 65 nm) and N_Bac_ = 71 docked vesicles / 57 docked vesicles (< 65 nm).

**Fig. 5.**
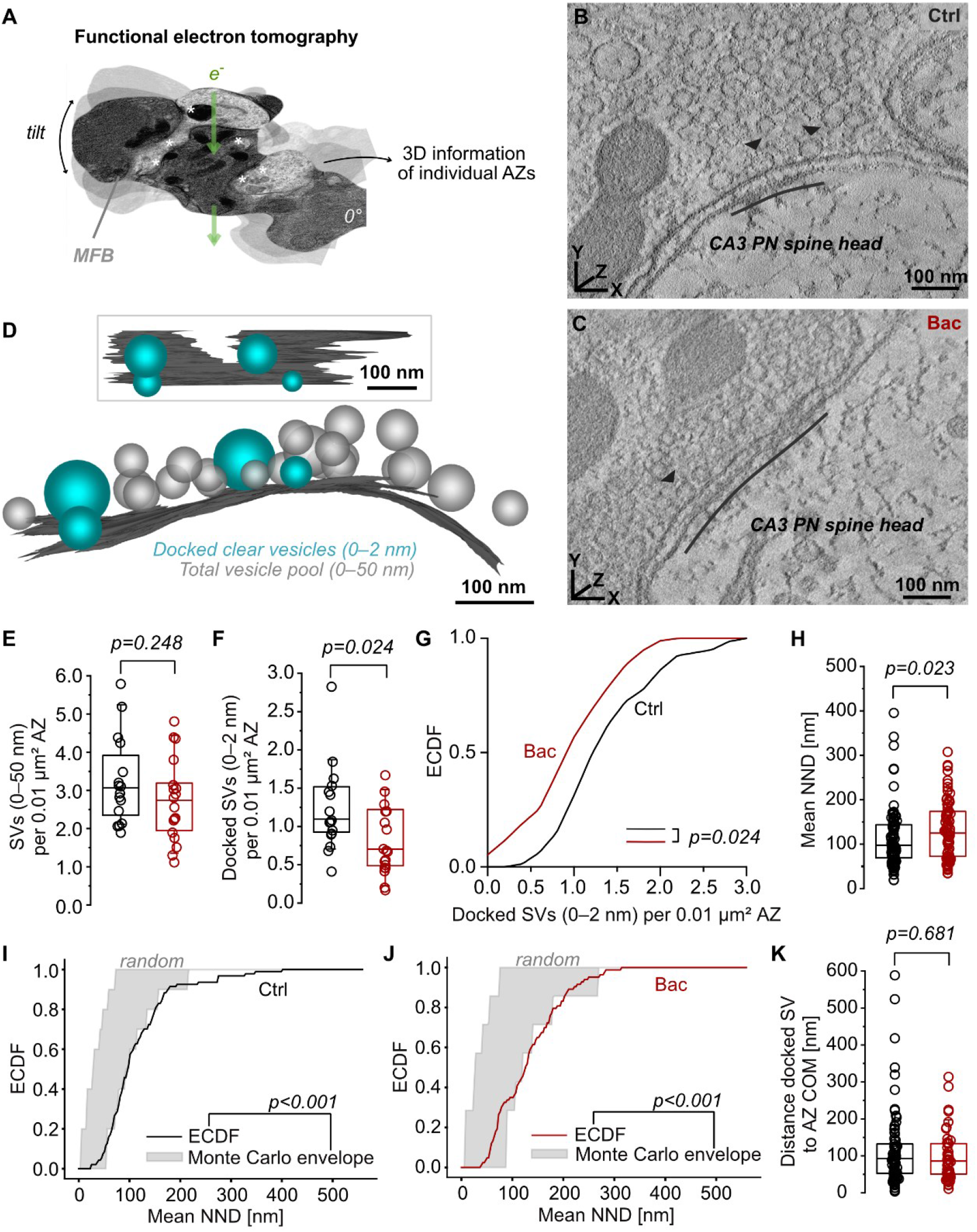
Presynaptic GABA_B_ receptor activation disperses the docked vesicle pool. **(A)** Schematic of the electron tomography (ET) workflow using high-pressure frozen acute brain slices (*35*). On an exemplary EM micrograph from a tilt series, asterisks mark postsynaptic sites positioned opposite to putative AZs, representing acquisition sites for 3D ET. **(B, C)** Representative electron tomographic slices of MFB AZs in Ctrl **(B)** and Bac **(C)** opposite to the postsynaptic density (black line), with docked SVs (0–2 nm; arrowheads). **(D)** 3D model of the AZ surface (dark gray) presented in **(C)**, containing total SV pool (0–50 nm; membrane-proximal pool (*3*); light gray), and docked clear SVs (0–2 nm; turquoise). Inset (top) depicts the orthogonal view. **(E, F)** Summary box plots of the number of total SVs (0–50 nm) and of docked clear SVs per 0.01 µm² AZ surface area in Ctrl and Bac. **(G)** ECDF plots of the data displayed in **(F).** Data in A–G from N_Ctrl_ = 16 tomograms (2 mice) and N_Bac_ = 19 tomograms (3 mice). **(H)** Summary box plot of the mean nearest neighbor distance (NND) for individual docked SVs in Ctrl and Bac. Only AZs with > 1 docked SVs were included. **(I, J)** ECDF plots of observed NNDs (solid line) overlaid on the 95% Monte Carlo envelope for Ctrl and Bac. ECDFs within the Monte Carlo envelope indicate complete spatial randomness; deviations above (shorter NNDs) indicate clustering, while below (longer NNDs) indicate spatial regularity (*89*), as observed here. **(K)** Summary box plot of the distance of docked clear SVs to the 3D AZ center of mass (COM) in Ctrl (N_Ctrl_ = 81 vesicles) and Bac (N_Bac_ = 48 vesicles). Data in H–K from N_Ctrl_ = 94 NNDs and N_Bac_ = 83 NNDs.

To obtain near-to-native synaptic ultrastructure, we combined drug incubation (baclofen, 5 min) and high-pressure freezing (“pharm-and-freeze”) of acute brain slices at physiological temperature (37°C) with two-dimensional (2D) transmission electron microscopy (EM) and 3D electron tomography (ET) analysis of the same sample (Fig. S5; Fig. 5A). 2D EM analysis revealed that the number of docked vesicles, defined as directly attached to the presynaptic membrane within 0–2 nm vertical distance (*35*, *53*), was significantly reduced (Fig. 4, A to D; Ctrl: median 0.43 (0.31–0.58, 25^th^–75^th^ percentile) vs. baclofen: 0.25 (0–0.45) docked vesicles per 100 nm AZ profile length; *p* < 0.001; 65 and 62 AZs, respectively). Additionally, we tested for the effect of baclofen on the docked vesicle diameter using 2D EM (Fig. 4, E to G), extending the docking criterion from 0–2 nm to 0–5 nm (*4*, *26*). Unexpectedly, this analysis revealed a significant diameter reduction under baclofen (baclofen vs. Ctrl: *p* = 0.040; Fig. 4E). Analysis of cumulative distributions indicated that the abundance of vesicles with larger diameter (‘giant vesicles’ (*55*), > 65 nm) appeared to be reduced by baclofen (Fig. 4, F to G; Table S6). These findings demonstrate that GABA_B_ receptor activation both reduces the docked vesicle pool and changes its composition. In summary, our results support the conclusion that inhibitory neuromodulation reduces RRP and docked vesicle pool in parallel, establishing a tight structure-function relation between functional and structural vesicle pools in MFBs.

### Presynaptic GABA_B_ receptor activation spatially disperses the docked vesicle pool at individual mossy fiber AZs

Previous results suggested that cAMP-mediated plasticity not only leads to a size increase of the docked vesicle pool but also to its spatial reconfiguration within AZs of MFBs (*4*, *25*, *56*, *57*). Does presynaptic inhibition revert or delete such structural remodeling? To further analyze the pool ultrastructure at the level of individual vesicles, we compared AZ ultrastructure using standardized 3D ET of near-to-natively high-pressure frozen, baclofen incubated and control samples previously analyzed by 2D EM (Fig. 5, A to D; Fig. S5 and Fig. S6).

Quantitative analysis of 3D AZ models confirmed the reduction of the docked clear vesicle pool (Fig. 5, E and F; Fig. S5; 0–2 nm; Ctrl: 1.10 (0.92–1.54) vs. baclofen: 0.70 (0.49–1.22) vesicles per 0.01 µm² AZ surface area, *p* = 0.024; Ctrl: 16 vs. baclofen: 19 AZ tomograms). This ultrastructural alteration was confined to the docked pool, as both the size of the membrane-proximal pool, encompassing one vesicle diameter plus a vesicle protein decoration zone (0–50 nm; Fig. 5D; Table S6; (*3*, *35*)), and the AZ surface area across conditions remained unchanged (Fig. S6; Table S6). To test the unexpected reduction in docked vesicle diameter observed in 2D, we analyzed 3D reconstructions. Docked clear vesicles (0–2 nm) did not differ in diameter between baclofen and control (*p* = 0.917; Fig. S6), but more loosely docked clear vesicles (2–5 nm) were smaller under baclofen (*p* = 0.009; Fig. S6). In addition, the total vesicle volume per AZ surface area was significantly reduced (*p* = 0.004; Fig. S5). In MFBs, the vesicle pool not only consists of clear vesicles but also of dense-core vesicles, whose exact contribution to synaptic release remains enigmatic (*3*, *35*, *58*). Under baclofen, docked dense-core vesicles had a significantly smaller diameter in baclofen (Fig. S6; *p* = 0.004), whereas their number and volume per AZ surface area were not significantly changed (Fig. S6; Table S6). Taken together, 3D ultrastructural analysis revealed that GABA_B_ receptor activation selectively reduces vesicle density and single-vesicle volume.

Based on the concept that AZ proteins compact during cAMP-induced potentiation at giant MFBs (*25*), we hypothesized that GABA_B_ receptor-mediated inhibition induces the opposite effect, spatial dispersion of AZ ultrastructure. Baclofen treatment significantly increased the nearest neighbor distance (NND; mean of the five closest vesicles) among docked clear vesicles to 124.8 nm (72.4–173.7 nm, 83 vesicles) compared to 97.4 nm (69.0–144.0 nm, 94 vesicles) in Ctrl conditions (Fig. 5G; *p* = 0.023), whereas the membrane-proximal pool remained unchanged (Table S6; *p* = 0.395). Spatial point pattern analysis using Monte Carlo envelopes (N = 999 independent simulations), controlling for vesicle density and AZ area as covariates, confirmed a non-random, spatially regular arrangement of docked vesicles in both conditions (Fig. 5, I and J; Table S6; *p* < 0.001). To further assess 3D vesicle positioning, we computed the distance from each docked vesicle to the AZ center of mass as a standardized localization metric (*35*). This analysis showed no significant difference between conditions (Fig. 5K), indicating no global spatial dispersion of membrane-proximal vesicles across the entire AZ. Instead, GABA_B_ receptor-mediated inhibition locally increased spacing between individual docked vesicles and their immediate neighbors at individual docking sites within the AZ. Thus, presynaptic inhibition reorganizes AZ ultrastructure by reducing the number and volume of docked vesicles, altering vesicle pool composition, and increasing the spatial dispersion of docked vesicles.

### Interneuron terminals adjacent to mossy fiber synapses are positioned for GABA spillover in CA3 *stratum lucidum*

Building on the observed functional and structural changes at individual MFBs following pharmacological activation of presynaptic GABA_B_ receptors, we next examined whether synaptically released GABA elicits comparable effects at hippocampal mossy fiber synapses under more physiological conditions. However, previous EM work on mossy-fiber associated interneurons did not report evidence for GABAergic synapses onto MFBs (*2*, *3*, *59*), leaving the mesoscale anatomy of such regulation unclear. Thus, we first quantified the 3D organization of GABAergic terminals relative to MFBs using light-microscopy-based connectomics (LICONN) analysis with iterative ∼16-fold tissue expansion (*60*) (Fig. 6; Fig. S7). Putative MFB AZs and interneuron terminals in CA3 *stratum lucidum* were identified by pan-protein staining combined with immunolabeling for Munc13-1 (a proxy for vesicle priming sites) and VGAT (vesicular GABA transporter, marking GABAergic terminals; Fig. 6, A to C). NND measurements between manually segmented MFBs and semi-automatically detected VGAT clusters (Fig. 6D) revealed a mean of 82 ± 5 VGAT clusters within a 5 µm radius of MFB outer membranes (Fig. 6E; 30 MFBs from 6 image volumes, 3 mice). VGAT clusters consisted of median 14.0 (4.0–89.0) VGAT punctae per cluster. The mean distance from an individual MFB to its nearest VGAT cluster was 0.30 ± 0.04 µm (Fig. 6F). Although direct synaptic contacts were not identified, subsets of inhibitory terminals were in close proximity to AZs in MFBs, suggesting that synaptically released GABA likely reaches presynaptic GABA_B_ receptors at mossy fiber synapses.

**Fig. 6.**
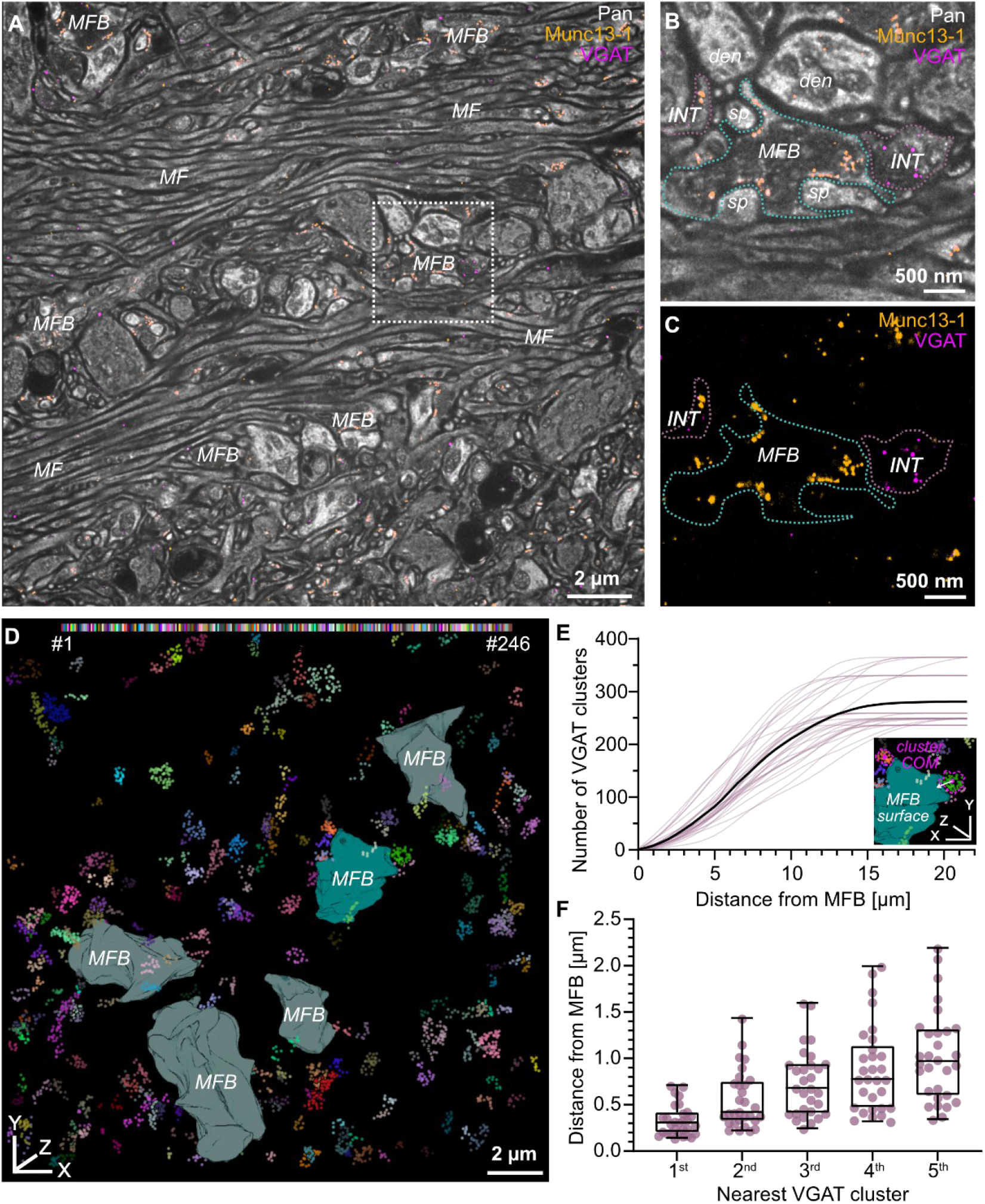
Expansion microscopy quantifies putative interneuron terminals in close proximity to MFBs in CA3 *stratum lucidum.* **(A–C)** Spinning-disc confocal micrographs of a 50-µm-thick hippocampal slice expanded ∼16-fold using LICONN iterative expansion microscopy combined with pan-protein staining and immunolabeling of Munc13-1 (AZs; yellow) and VGAT (putative interneuron terminals (INTs); magenta). White dotted box in overview **(A)** indicates the location of the region of interest in **(B, C)**. The putative MFB and INTs were outlined in cyan and dusted pink dotted lines, respectively. Scale bars reflect pre-expansion lengths. Abbreviations: MF, mossy fibers; den, dendrite; sp, spine head. **(D)** Manually segmented MFBs with semi-automatically detected VGAT clusters using Density-Based Spatial Clustering of Applications with Noise (DBSCAN) in a 20.3 µm × 20.3 µm × 10.9 µm large image volume of CA3 *stratum lucidum*. The reconstructed MFB in cyan corresponds to the MFB outlined in **(B, C)**. Individual VGAT clusters are randomly color-coded for visual clarity. **(E)** Cumulative plot of the number of VGAT clusters against distance from individual MFBs (dusted pink), overlaid with the average number of VGAT clusters (black). Inset shows a schematic of the distance measurement between the 3D COM of a cluster and the nearest surface of a segmented MFB. **(F)** The summary box plots of the distances of the five nearest VGAT clusters from individual MFBs. Data in E andF from N = 30 MFBs (3 mice).

### Synaptic activation of GABA_B_ receptors suppresses mossy fiber transmission and limits plasticity in the structural-functional readily releasable pool

Based on both the quantified mesoscale anatomy of inhibitory terminals surrounding MFBs and previously reported CA3 interneuron activity from *in vivo* recordings (*61–63*), we employed an optogenetic approach to examine presynaptic inhibition induced by synaptically released GABA. To achieve Cre-dependent channelrhodopsin (ChR2) expression exclusively in GABAergic interneurons, we crossed a Vgat-Cre mouse line with an Ai32 reporter line [Ai32(RCL-ChR2(H134R)/EYFP)]. By restricting the illumination area to a confined area in CA3 *stratum lucidum*, inhibitory axonal arborizations (*59*) were optogenetically activated in close proximity to the individually measured mossy fiber bouton with blue light pulses (Fig. 7A).

**Fig. 7.**
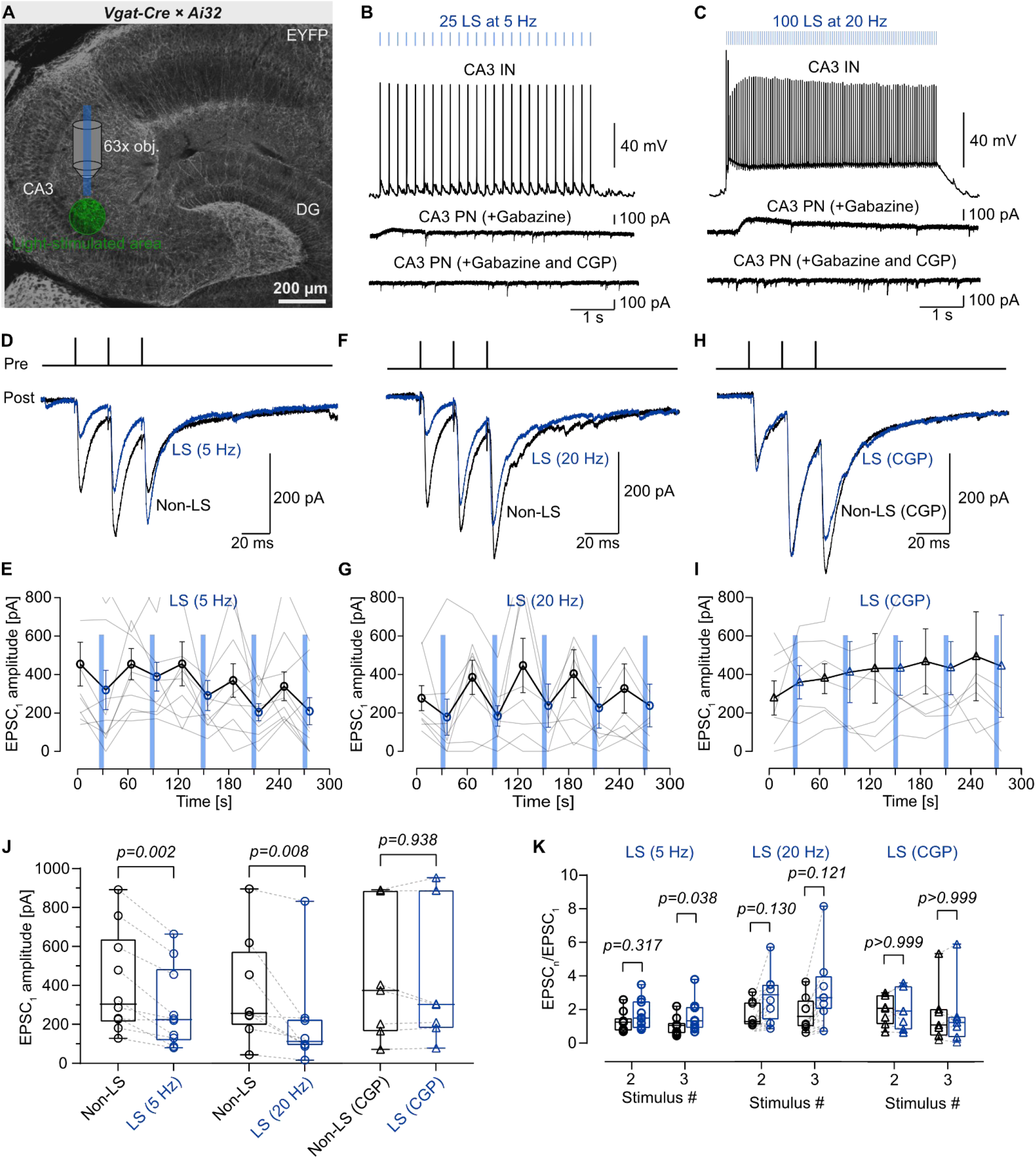
Synaptically released GABA from optogenetically stimulated CA3 *stratum lucidum* interneurons suppresses synaptic transmission at MFB–CA3 PN synapses. **(A)** Maximum intensity Z-projection of a confocal micrograph of a 50-µm-thick hippocampal slice from a Vgat-ires-Cre x Ai32 (RCL-ChR2(H134R)/EYFP) mouse, showing ChR2-EYFP expression in VGAT^+^ interneurons (INs). The dentate gyrus (DG) and CA3 region are annotated. Light stimuli were applied to a confined area in CA3 through a 63x objective (obj.). **(B, C)** Representative traces of evoked APs in a CA3 INs (top) and outward K^+^ currents in CA3 PNs (middle) during a 5-Hz train of 25 light stimuli (LS) and 20-Hz train of 100 LS. Outward K^+^ currents in CA3 PNs were blocked in the presence of 5 µM GABA_B_R antagonist CGP 55845 (CGP; bottom). Light intensity: 5 mW mm^-2^, duration: 5 ms. **(D, F, H)** Representative traces of EPSCs evoked by a 50-Hz train of 3 stimuli without light stimulation (Non-LS; black) and 1 s after a train of LS (blue) at 5 Hz (LS (5 Hz)), 20 Hz (LS (20 Hz)), or 20 Hz in the presence of CGP (LS (CGP)). **(E, G, I)** EPSC_1_ peak amplitudes plotted against experimental time for protocols alternating periods of Non-LS and LS (5 Hz), LS (20 Hz), and LS (CGP). Light blue shadings indicate LS periods. **(J, K)** Summary box plots of EPSC_1_ peak amplitudes and EPSC_n_/EPSC_1_ for stimulus number 2 and 3 in Non-LS versus LS (5 Hz) Non-LS versus LS (20 Hz), and Non-LS versus LS (CGP). MFB–CA3 PN pairs: N = 10 for LS (5 Hz), N = 8 for LS (20 Hz), and N = 7 for LS (CGP).

We performed paired pre-postsynaptic recordings and compared EPSC_1_ amplitudes with and without light stimulation at two physiologically relevant interneuron firing frequencies. We started with 25 light stimuli (LS) at 5 Hz, resembling a firing pattern of mossy fiber associated, likely CCK-expressing interneurons (*14*, *59*, *61*), and additionally tested 100 LS at 20 Hz, resembling parvalbumin-positive interneurons (*62*) and representing the maximum frequency at which reliable ChR2 stimulation could be achieved ((*26*), Fig. 7, B and C). For both frequencies, 5-s light stimulation evoked slow inhibitory postsynaptic currents (IPSCs) in CA3 PNs, which could be blocked by the selective GABA_B_ receptor antagonist CGP 55845 (5 µM), confirming GABA_B_ receptor-mediated activation of postsynaptic GIRK channels (Fig. 7, B and C). Both light stimulation protocols significantly reduced the amplitude of EPSC_1_ compared to interleaved Ctrl (Fig. 7, D to J; Non-LS 410.6 ± 82.7 pA; LS (5 Hz): 282.7 ± 65.5 pA, *p* = 0.002, 10 MFB–CA3 PN pairs; Non-LS 368.1 ± 97.3 pA; LS (20 Hz): 213.4 ± 91.8 pA, *p* = 0.008, 8 MFB−CA3 PN pairs). Consistent with the effects of baclofen, light stimulation increased the multiple-pulse ratio in a frequency-dependent manner, with the EPSC_3_/EPSC_1_ ratio significantly increased by 5 Hz (*p* = 0.038), while EPSC_2_ /EPSC_1_ and EPSC_3_ /EPSC_1_ ratios induced by 20 Hz showed a non-significant trend toward increase (Fig. 7K; Table S1). The effects of LS were blocked by CGP 55845, demonstrating the involvement of presynaptic GABA_B_ receptors (Fig. 7, H to K; Table S1; 7 MFB−CA3 PN pairs).

To assess the ultrastructural impact of synaptically released GABA on MF transmission, we performed flash-and-freeze experiments (*26*) with the described Vgat-Cre x Ai32 mice. At defined time points after application of LS equivalent to the paradigm used in electrophysiology, acute slices were subjected to high-pressure freezing and processed for 2D EM (Fig. 8, A to C). For 5 Hz, the number of docked vesicles at MFB AZs was significantly reduced (Fig. 8D; non-LS: median 0.66 (0.51–0.81) vs. LS (5 Hz): 0.31 (0–0.49) docked vesicles per 100 nm AZ profile length, *p* < 0.001, 61 and 85 AZs, respectively). A comparable reduction was observed at 20 Hz, both in the absence of gabazine (Fig. 8D; non-LS: median 0.60 (0.49–0.84) vs. LS (20 Hz): 0.30 (0.15–0.48) docked vesicles per 100 nm AZ profile length; *p* < 0.001; 72 and 84 AZs, respectively) and in its presence (Fig. 8D; non-LS: median 0.68 (0.53–0.84) vs. LS (20 Hz) + gabazine: 0.34 (0.21–0.43) docked vesicles per 100 nm AZ profile length, *p* < 0.001, 54 and 62 AZs, respectively). Together, these results establish that synaptically released GABA, specifically acting through presynaptic GABA_B_ receptors, functionally and structurally suppresses mossy fiber transmission at the level of individual boutons and their AZs.

**Fig. 8.**
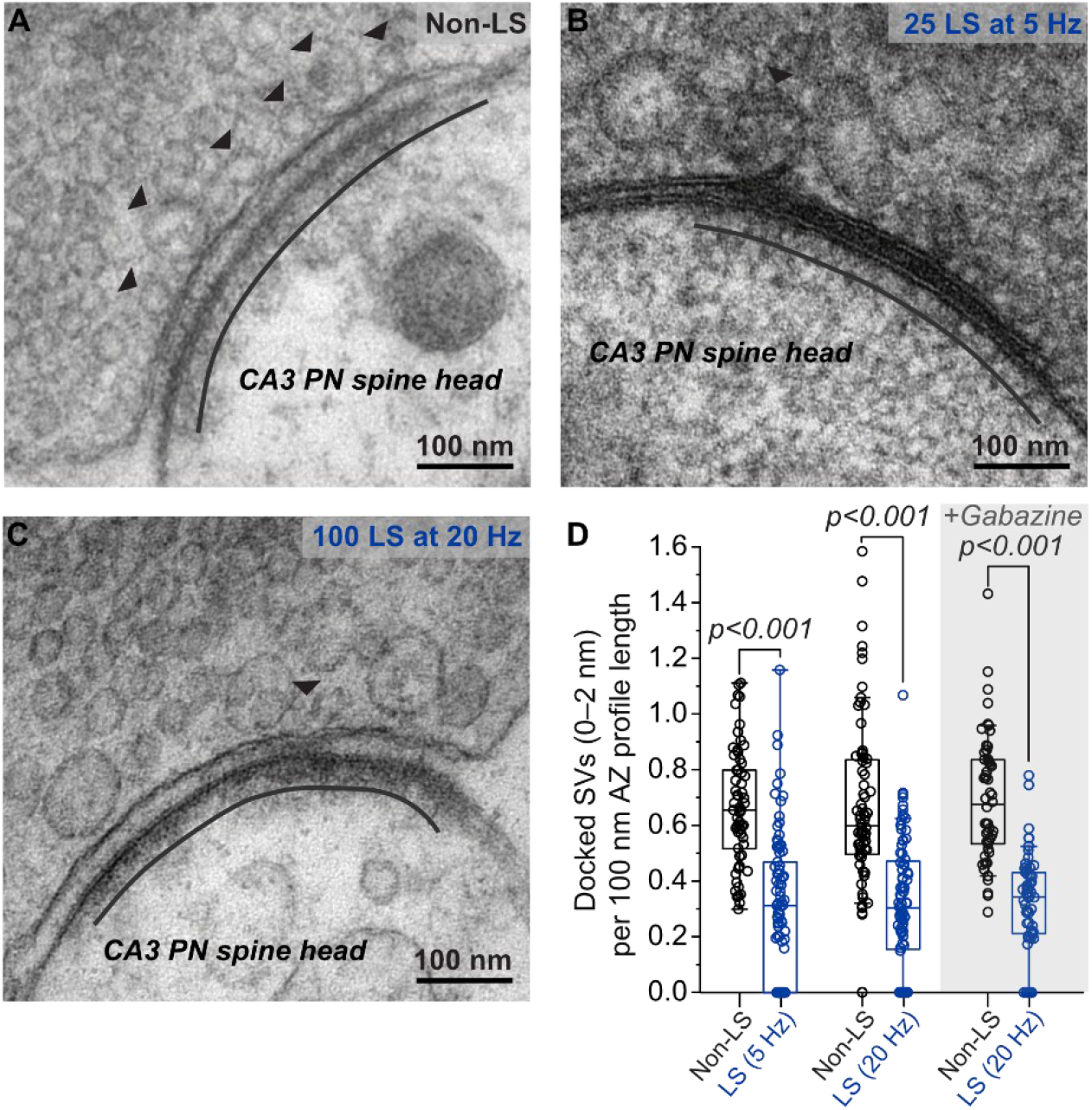
Synaptically released GABA reduces the docked vesicle pool at MFB–CA3 PN synapses. **(A–C)** Representative 2D electron micrographs of MFB AZs (70-nm ultrathin sections), defined as structural compartments with docked synaptic vesicles (SVs, arrowheads) opposite to the postsynaptic density (extent indicated by black lines), from Non-LS tissue **(A)** and ‘flash- and-freeze’ tissue with 25 LS at 5 Hz **(B)** or 100 LS at 20 Hz **(C)**. **(D)** Summary box plots of docked SV number (0–2 nm) per 100 nm AZ profile length in Non-LS versus LS (5 Hz), Non-LS versus LS (20 Hz), and Non-LS versus LS (20 Hz) in the presence of gabazine. Data in D from 5 Hz: N_Non-LS_ = 61 AZs (2 mice), N_5Hz_ = 85 AZs (2 mice); 20 Hz: N_Non-LS_ = 72 AZs (2 mice), N_20Hz_ = 84 AZs (2 mice); 20 Hz with gabazine: N_Non-LS_ = 54 AZs (2 mice), N_20Hz + gabazine_ = 62 AZs (2 mice).

In rats, mossy fiber synapses exhibit robust PTP following 100-Hz stimulation, driven primarily by an expansion of the RRP and the docked vesicle pool (*4*). In mice, PTP and the underlying mechanisms have not been investigated. To test whether synaptically released GABA modulates this form of plasticity via presynaptic inhibition in mice, we combined the same high-frequency stimulation with a 5-s light stimulation on *stratum lucidum* interneurons during paired recordings. Consistent with previous findings in rats (*4*), albeit with a smaller effect magnitude, a single train of 100 stimuli at 100 Hz (HFS_100APs_) reliably induced PTP at mouse mossy fiber synapses (195.8 ± 55.0%, τ = 77.0 s; Fig. 9, A, B, and G). Optogenetic activation of interneurons at 20, 40, and 60 s after HFS largely abolished PTP, as quantified by unitary EPSCs before and after induction (Fig. 9, C to G). Without light stimulation, HFS markedly increased the EPSC_1_ peak amplitude from 193.0 ± 72.0 pA (Ctrl) to 433.3 ± 98.5 pA (HFS) (Fig. 9H; *p* = 0.002; 10 MFB–CA3 PN pairs). In contrast, under application of LS at 5 Hz and 20 Hz, HFS produced only a modest increase in EPSC_1_ peak amplitude (Fig. 9H; Ctrl: 247.3 ± 50.4 pA, HFS + LS (5 Hz): 332.3 ± 89.0 pA, *p* = 0.109, 8 MFB–CA3 PN paired pairs; Ctrl: 251.6 ± 90.0 pA, HFS + LS (20 Hz): 307.0 ± 100.1 pA, *p* = 0.105, 10 MFB–CA3 PN paired pairs), and left paired- and multiple-pulse ratios largely unchanged (Fig. S8; Table S4).

**Fig. 9.**
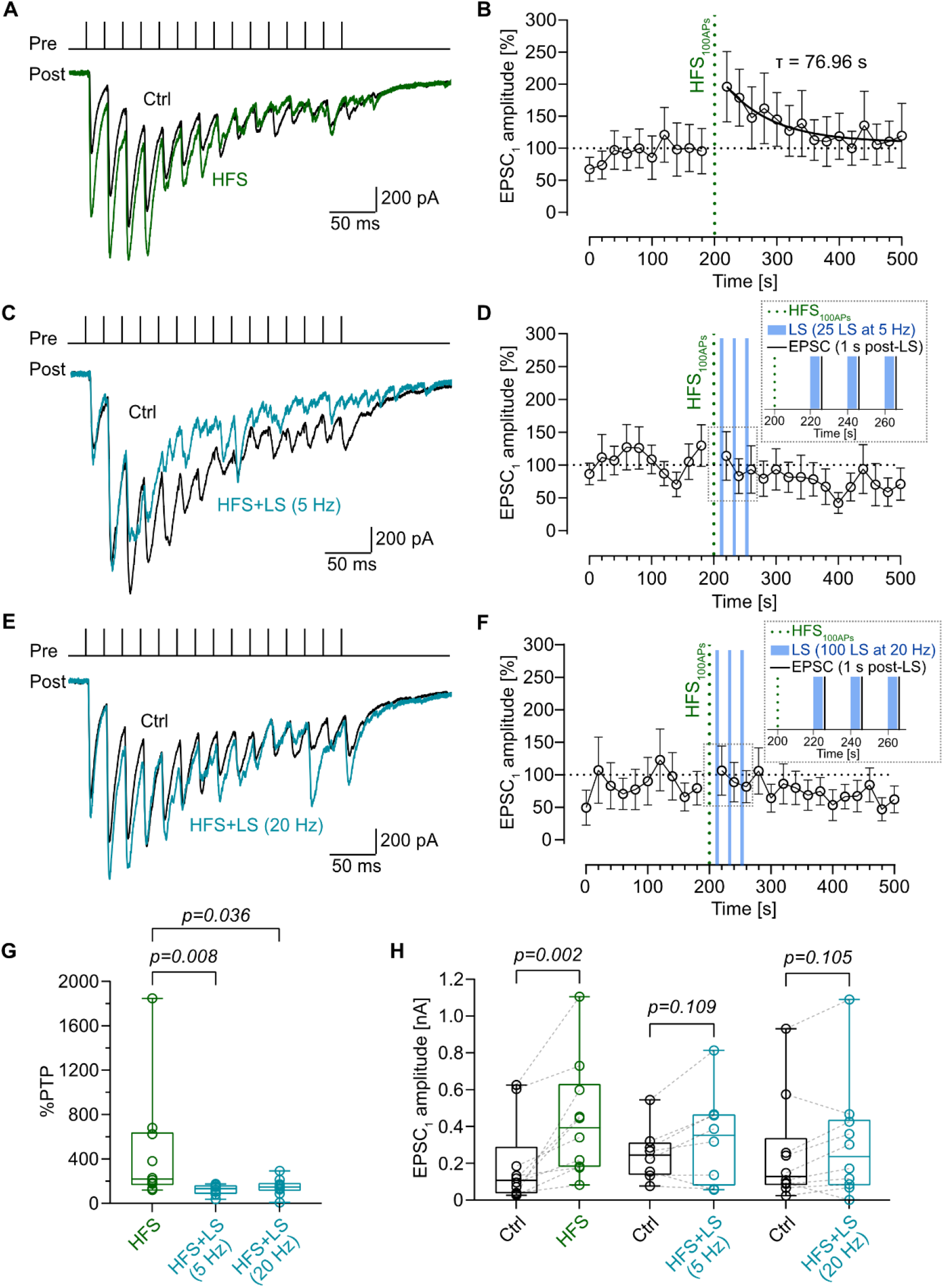
Synaptically released GABA limits post-tetanic potentiation at MFB–CA3 PN synapses. **(A)** Representative traces of EPSCs evoked by a 50-Hz train of 15 stimuli before HFS_100APs_ (Ctrl; black) and after HFS_100APs_ (HFS; green) from the same MFB–CA3 PN pair. **(B)** Normalized EPSC_1_ peak amplitudes (to the average EPSC_1_ peak amplitude from Ctrl) plotted against experimental time. The vertical dotted line indicates the time point of HFS_100APs_. **(C, E)** Representative traces of EPSCs evoked by a 50-Hz train of 15 stimuli before HFS_100APs_ (Ctrl; black) and after HFS_100APs_ followed by a 25 LS at 5 Hz (HFS (5 Hz); teal) or 100 LS at 20 Hz (HFS+LS (20 Hz)) from the same MFB–CA3 PN pair. **(D, F)** Normalized EPSC_1_ peak amplitudes plotted against experimental time for HFS+LS (5 Hz) and HFS+LS (20 Hz). The vertical dotted line indicates the time point of HFS_100APs_. Lightblue shading indicates LS periods. Insets show schematics of HFS_100APs_, LS, and EPSC recording at an expanded time scale. **(G, H)** Summary box plots of post-tetanic potentiation (PTP) magnitude (see Materials and Methods) and EPSC_1_ peak amplitudes in HFS, HFS+LS (5 Hz), and HFS+LS (20 Hz) with respective Ctrl. Data in B, D, F–H from N = 10 for HFS, 8 for HFS+LS (5Hz), and 10 for HFS+LS (20 Hz) MFB–CA3 PN pairs.

Consistent with previous observations (*4*), PTP in mice was associated with a significant increase in the RRP size based on cumulative EPSC analysis (Fig. 10, A and D; SMN method: Ctrl: 1.24 ± 0.27 nA; HFS: 2.02 ± 0.32 nA, *p* = 0.002; TR method: Fig. S8 and Table S4, *p* = 0.004), whereas light activation of interneurons kept the RRP size close to Ctrl (Fig. 10, B to D; Ctrl: 1.17 ± 0.19 nA, HFS + LS (5 Hz): 1.23 ± 0.30 nA, *p* = 0.641; Ctrl: 1.26 ± 0.32 nA, HFS + LS (20 Hz): 1.46 ± 0.27 nA, *p* = 0.160). HFS enhanced P_v_ and decelerated the vesicle refilling, whereas both effects were abolished by light-induced GABA release (Fig. 10, E and F; Table S4). Collectively, these results demonstrate that synaptically released GABA engages presynaptic GABA_B_ receptors to modulate short-term plasticity at the single-bouton level, inhibiting transmitter release and limiting memory storage in the readily releasable pool (Fig. 10G).

**Fig. 10.**
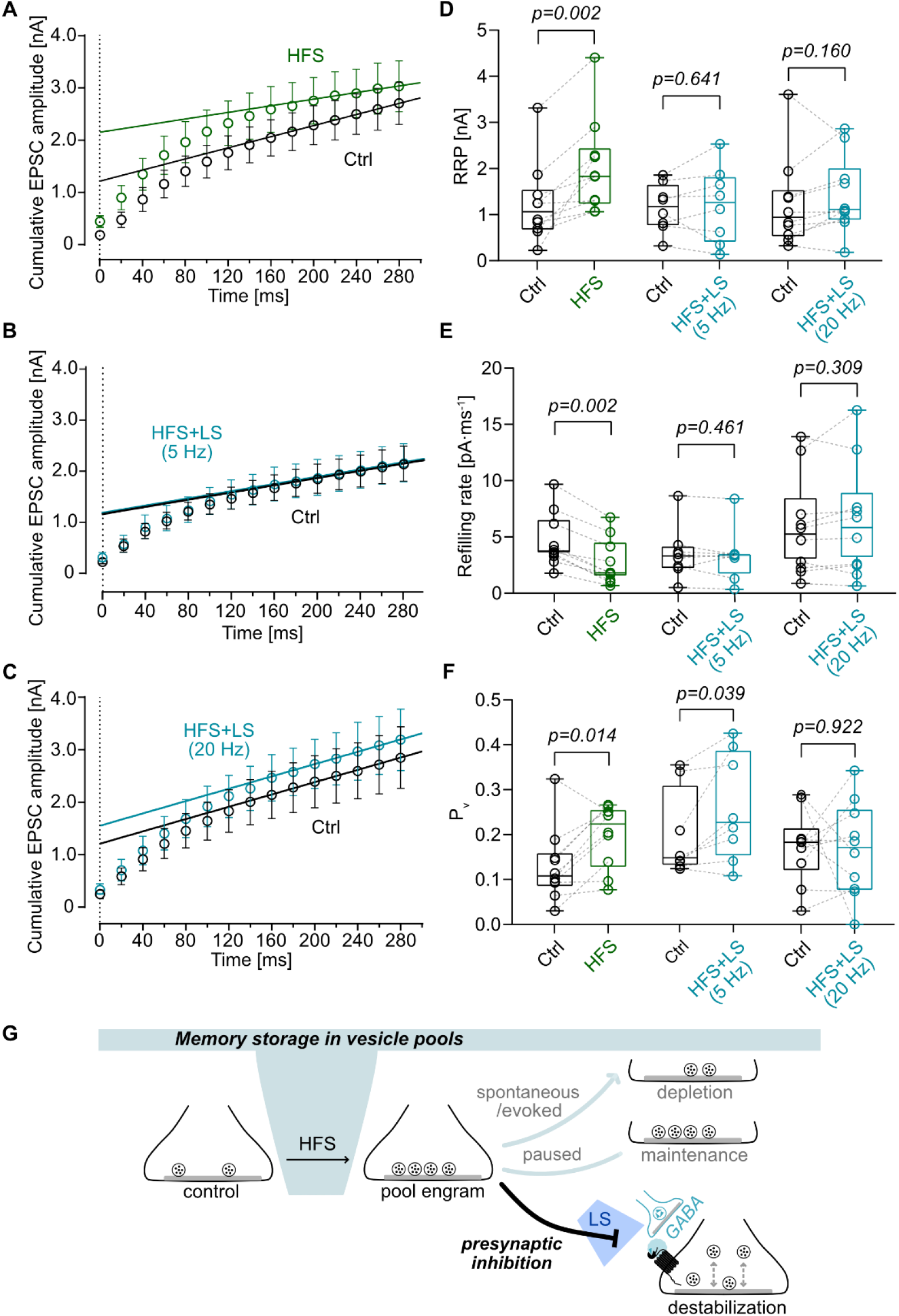
Synaptically released GABA modulates PTP-induced vesicle pool engram formation at MFB–CA3 PN synapses. **(A–C)** Cumulative plots of EPSC peak amplitudes evoked by a 50-Hz train of 15 stimuli (Ctrl) and after HFS, HFS+LS (5 Hz), and HFS+LS (20 Hz). RRP size, refilling rate, and P_v_ were estimated by linear regression of the last four data points and back-extrapolation to time 0. **(D–F)** Summary box plots of RRP size, refilling rate, and P_v_ in Ctrl versus HFS, Ctrl versus HFS+LS (5 Hz), and Ctrl versus HFS+LS (20 Hz). Data in A–F from N = 10 for HFS, 8 for HFS+LS (5 Hz), and 10 for HFS+LS (20 Hz) MFB–CA3 PN pairs. **(G)** Dynamics of the vesicular pool engram across activity states. In control conditions, high-frequency stimulation (HFS) expands the docked vesicle pool (left), forming a structural pool engram (middle). Spontaneous or evoked release depletes this engram (top right), whereas inactivity (“paused”) preserves it (middle right). Under presynaptic inhibition via synaptically released GABA, evoked by LS, the pool engram is reorganized through altered docking and undocking (bottom right).

## DISCUSSION

Although presynaptic inhibition is a well-established mechanism of synaptic regulation, its structural basis and its consequences for information storage in cortical circuits remain poorly understood. Our results provide evidence that presynaptic inhibition shapes hippocampal circuit operations in multiple ways. First, our biophysical analysis at hippocampal mossy fiber synapses reveals that GABA_B_ receptor activation by the specific agonist baclofen reduces release probability via a reduction of the presynaptic Ca^2+^ current (*20*), but in addition shrinks the RRP, revealing a second, mechanistically distinct arm of presynaptic inhibition. Thus, our results add to the emerging evidence that the RRP is a key target of synaptic neuromodulation (*4*, *23*, *25*, *64*). Second, our structural analysis by near-to-native 2D-EM and 3D-ET reveals that GABA_B_ receptor activation reduces the size of the docked synaptic vesicle pool and alters its nanoscopic configuration, identifying a major structural correlate of presynaptic inhibition. Third, synaptically released GABA reduces transmitter release, docked vesicle pool size, and presynaptic plasticity. These effects are most likely facilitated by the close proximity of GABAergic terminals in *stratum lucidum* and presynaptic GABA_B_ receptors in MFBs. Thus, GABA_B_ receptor-mediated inhibition limits the persistence of vesicle pool plasticity at MF synapses of hippocampal CA3 (*4*).

### Mechanisms of presynaptic inhibition

It has long been assumed that presynaptic GABA_B_ receptor-mediated inhibition reduces release probability primarily by decreasing presynaptic Ca^2+^ inflow (*20*). However, our results show that both changes in Ca^2+^ inflow and RRP are involved (Fig. 3). Previous work suggested that modulation of presynaptic Ca^2+^ channels by G-protein coupled receptors is mediated by direct binding of ꞵɣ subunits to the channel’s ɑ subunit (*18*, *19*). ꞵɣ subunits selectively inhibit voltage sensor-domains I and IV, which induces reluctant gating (*65*, *66*) and thereby limits Ca^2+^ influx during short presynaptic APs (*67*). Consistent with this, baclofen reduced the Ca^2+^ charge (∼40%) proportionally more than the peak current (∼35%) (Fig. 3, Table S5). Further, MFBs contain a mixture of P/Q-, N- and R-type Ca^2+^ channels (*68*, *69*). Previous work suggested N-type Ca^2+^ channels as primary neuromodulatory targets (*70*, *71*). Our results are consistent with this view, since inhibition of presynaptic Ca^2+^ current by baclofen (∼35%) closely matches the proportional contribution of N-type Ca^2+^ channels to the total presynaptic Ca^2+^ current (∼27%) in hippocampal MFBs (*69*).

By contrast, regulation of the vesicle pool involves the G-protein ɑ subunits and the cAMP pathway. Early studies on GABA_B_ receptor activation suggested that baclofen inhibits adenylate cyclase (AC) activity (*27*). Furthermore, recent work indicates that AC, cAMP, and cAMP-dependent protein kinase A (PKA) regulate the size and configuration of vesicular pools (*25*, *23*). Finally, several PKA targets expressed in presynaptic terminals are implicated in the regulation of vesicular pools (*72*). For example, phosphorylation of synapsin may modulate its interaction with actin and thereby alter the organization of the synaptic vesicle pool (*73*).

Why are two pathways needed to control transmitter release from MFBs, one targeting Ca^2+^ channels and another regulating the RRP (Fig. 3I)? One potential reason might be amplification (Table S5). Because of the power relationship between Ca^2+^ inflow and transmitter release, a 35% reduction in Ca^2+^ current translates into a larger change in P_v_. With a power coefficient of 1.77 at this synapse (*52*), this predicts a 50% reduction of P_v_, closely replicating the measured decrease (Fig. 3). The decrease in P_v_ then combines multiplicatively with the even greater reduction in RRP. Given that release = P_v_ × RRP, the predicted total reduction of 87% closely matches the measured value (Table S5). A second potential reason lies in the differential kinetics of the two pathways. Whereas the ꞵɣ-subunit-mediated effects on Ca^2+^ channels occur within seconds, cAMP/PKA-dependent regulation of pool organization operates on a time scale of tens of seconds or longer (*23*, *25*, *72*). Thus, GABA_B_ receptor-mediated presynaptic inhibition integrates rapid suppression of Ca^2+^ channel gating with a slower modulation of vesicle pool size, thereby providing an exquisitely broad kinetic range of presynaptic control.

### Structural and functional changes during presynaptic inhibition

Our ultrastructural analysis reveals that activation of GABA_B_ receptors by baclofen markedly reduced the size of the docked vesicle pool at AZs of hippocampal MFBs (Fig. 4; Fig. 5). Importantly, synaptically released GABA reproduced the reduction in the docked vesicle pool (Fig. 8), indicating that this structural effect is not restricted to saturating pharmacological activation. Together with our functional data demonstrating that GABA_B_ receptor activation also reduces the RRP size (Fig. 3; Fig. 10; Fig. S4; Fig. S8), these results indicate that presynaptic inhibition regulates both the functional and structural vesicle pools in parallel. Earlier studies found the same parallel regulation in the opposite direction: cAMP-mediated activation by forskolin increased both docked pool and RRP (*25*, *57*). Thus, the docked vesicle pool and the RRP likely overlap substantially.

However, previous studies have consistently reported that the RRP is smaller than the docked vesicle pool (*25*, *53*, *74*). Our results provide a framework to reconcile this apparent discrepancy: the RRP may correspond to a specific subset of docked vesicles. Two observations support this idea: First, the nanometer resolution EM tomography allowed us to subdivide docked vesicles by their distance to the AZ membrane. Baclofen selectively reduced the pool of tightly docked clear vesicles (0–2 nm from the AZ membrane), whereas loosely docked vesicles (2–5 nm (*35*, *74*)) and the membrane-proximal vesicle pool (0–50 nm (*3*)) as a whole did not change significantly (Fig. S6). Second, the tightly docked pool is ∼3-fold smaller than the membrane-proximal vesicle pool (Fig. 5), providing closer agreement with the RRP. Accordingly, the tightly docked vesicle pool represents a closer structural correlate of the RRP than the loosely docked or other membrane-proximal pools.

The nanoscale spatial organization of signaling events and their structural correlates is a critical determinant of synaptic efficacy and dynamics (*7*, *52*, *75*). Our 3D analysis demonstrates that GABA_B_ receptor activation increases the spatial dispersion of the docked vesicle pool (Fig. 5). This dispersion was not random: docked vesicles adopted a regular, non-clustered arrangement on the AZ membrane (Fig. 5), consistent with a higher-order spatial organization of the AZ. Such a dispersion may increase the coupling distance between presynaptic Ca^2+^ channels and release sensors. In line with looser coupling, baclofen slightly but significantly increases EPSC_1_ latency (Fig. 1). Vesicle dispersion has also been implicated in plasticity. Recent work proposed that a dispersion of the total vesicle pool in boutons may contribute to LTP expression (*57*, *76*). Further work linking AZ nanostructure to synaptic function will be needed to clarify the relevance of such spatial vesicle patterns for synaptic transmission and plasticity.

Beyond number and position, GABA_B_ receptor activation also affects which vesicles remain docked. In comparison to other synapses, hippocampal mossy fiber synapses show a wide distribution of clear vesicle diameters (*35*, *55*, *77*). In single 2D sections, baclofen reduced the diameter of docked vesicles (0–5 nm to the AZ membrane) and the proportion of large clear vesicles (Fig. 4). 3D tomography, which provides reliable size estimates (*35*, *53*, *55*), revealed that the diameter reduction was confined to loosely docked clear vesicles (2–5 nm; Fig. S6). Together with the reduced mEPSC amplitude (< 1 ms; Fig. S2) under baclofen, this suggests that loosely docked vesicles contribute to spontaneous release and that their smaller size underlies the mEPSC amplitude reduction. The size reduction could reflect size-dependent selection during priming or a dynamic change in vesicle size: In kiss-shrink-run-like fusion, vesicles release transmitter through a transient pore, shrink, and detach intact (*78*, *79*), a mode that Gᵢ/ₒ-coupled receptors may favor via G_βγ_ (*80*), and may enrich shrunken vesicles near the AZ. Dense-core vesicles, potentially containing brain-derived neurotrophic factor (*58*), also changed in size, but, unlike the docked clear pool, not in docking density (Fig. S6). Further work will be needed to distinguish the contribution of vesicle ‘sub-pools’, types, and release modes to transmitter release at mossy fiber synapses.

### Physiological significance of presynaptic inhibition

Our results demonstrate that presynaptic inhibition of mossy fiber synaptic transmission is not only initiated by baclofen, but also by synaptically released GABA (Figs. 7–10). Thus, presynaptic inhibition may represent a major contributor to inhibition in the CA3 circuit (*61–63*). Although the exact cellular source of the synaptically released GABA remains unidentified, our data support the idea that neighboring interneuron terminals may play a critical role (Fig. 6). Likely candidates include CCK-positive mossy fiber-associated interneurons (*59*, *81*) or NOS-positive Ivy cells, whose axons arborize in *stratum lucidum*. Because the somatodendritic domains of these interneurons are also confined to this layer, they may form a feedforward inhibition circuit with a previously unrecognized presynaptic component (*82*), characterized by a kinetically delayed but more persistent inhibitory effect.

Our results suggest that the GABA_B_ receptor-mediated inhibition substantially changes single-synapse computations (*36*, *37*). Under control conditions, the mossy fiber synapse has been depicted as a conditional detonator, in which a single synaptic event in conjunction with facilitation, potentiation, or neuromodulation can elicit APs in postsynaptic CA3 PNs (*41*, *42*). Hence, mossy fiber inputs function as an “OR gate”, whereby activation of any single input is sufficient to drive postsynaptic activity. After GABA_B_ receptor activation, the synapse is switched into a subdetonation mode (Fig. 2). Thus, activation of multiple converging inputs is required to activate the postsynaptic cell, consistent with an “AND gate” mechanism (Fig. 2F). This may enhance pattern separation at the synaptic mossy fiber–CA3 station along the trisynaptic circuit (*83*).

Mossy fiber synapses, through their presynaptic forms of plasticity, have been proposed to store information as vesicle pool engrams, reminiscent of a ‘nanosubstrate’ of the classical engram (*4*, *29*). Our results reveal how presynaptic inhibition may act on such a substrate. Both baclofen and synaptically released GABA reduced the tightly docked vesicle pool (Fig. 4; Fig. 5; Fig. 8), the very pool whose expansion constitutes the vesicle pool engram. Within this concept, GABA_B_ receptor-mediated presynaptic inhibition could negatively calibrate vesicle pool engrams.

Does presynaptic inhibition act on induction, readout, or maintenance of such engrams? Induction is unlikely to be involved, since light stimulation followed HFS by at least 20 s. If presynaptic inhibition would transiently suppress the readout, one would expect a joint reduction of P_v_ (via the fast arm, potentially mediated by ꞵɣ subunits) and RRP (via the slow arm, presumably involving cAMP). However, this was not observed (Fig. 10F). Furthermore, if synaptic GABA release only influenced the readout, the enlarged pool would be spared rather than consumed. Thus, the next electrical stimulation of MFBs after light stimulation should reveal an overshoot, analogous to the rebound observed after a stimulation pause following PTP induction (*4*). However, this was not observed (Fig. 9 D, F). Together, these findings suggest that GABA release interferes with the maintenance of the vesicle pool engram. Here, presynaptic inhibition may attenuate or even delete vesicle pool engrams. Future experiments at both subcellular and behavioral level will be needed to further test this provocative hypothesis.

The present findings have important implications for understanding brain diseases. In mice, deletion of GABA_B1a_, the principal subunit isoform at MFBs (*16*), causes epilepsy and memory deficits (*84*, *85*). The epilepsy phenotype is consistent with the loss of a brake on a conditional detonator synapse, whereas the memory deficits have been attributed to saturation of synaptic plasticity. In humans, loss-of-function variants in adherens junction–associated protein 1, which recruits GABA_B1a_ containing receptors to presynaptic sites, cause developmental delay, intellectual disability, and epilepsy – the first case of selective presynaptic GABA_B_ dysfunction (*86*). As mossy fiber synapses are highly enriched in *puncta adherentia* (*2*, *3*), deficits in mossy fiber transmission may substantially contribute to this phenotype. Impaired presynaptic inhibition of mossy fiber transmission and plasticity may therefore represent a common mechanism underlying hippocampal dysfunction across brain diseases.

## MATERIALS AND METHODS

### Animals

Animals were housed and bred in the Preclinical Facility at ISTA. All procedures were performed in strict accordance with institutional, national, and European guidelines for animal experiments, approved by the Bundesministerium für Bildung, Wissenschaft und Forschung of Austria (Reference numbers 2020–0.648.587 and 2022–0.121.440). For optogenetic manipulation of interneurons in patch-clamp recordings, a Vgat-ires-Cre line (RRID: IMSR_JAX:028862) was crossed with an Ai32 reporter line (RCL-ChR2(H134R)/EYFP, RRID: IMSR_JAX:012569, all available from Jackson Laboratory). For a granule cell specific knockout of GABA_B_1 receptors, a GABA_B_1 fl/fl line (provided by B. Bettler, (*40*)) was crossed with a Prox1-Cre line (RRID: MMRRC_036644-UCD, (*26*)).

### Acute hippocampal slice preparation

Acute hippocampal slices from 20-to 25-day-old (P20–25) C57BL/6J mice (RRID: IMSR_JAX:000664) were used for patch-clamp recordings and for ultrastructural analysis. Additionally, acute hippocampal slices from Vgat-ires-Cre x Ai32 mice were used for patch-clamp recordings. Slices of either 300 μm (patch-clamp recordings) or 150/200 μm (ultrastructural analysis) thickness were prepared according to the previously described protocols (*34*). After brief anesthesia with isoflurane and decapitation, brains were rapidly extracted from the skull and prepared for vibratome cutting. Slices were cut from either the left and/or right hemisphere using ‘magic cut’ mounting in ice-cold high-sucrose-based artificial cerebrospinal fluid (ACSF) with a vibratome (VT1200, Leica Microsystems). High-sucrose-based ACSF contained 64 mM NaCl, 120 mM sucrose, 25 mM NaHCO_3_, 10 mM D-glucose, 2.5 mM KCl, 1.25 mM NaH_2_PO_4_, 0.5 mM CaCl_2_, and 7 mM MgCl_2_, equilibrated with 5% CO_2_/95% O_2_, with an osmolarity of ∼334 mOsm. Slices were then recovered in the same high-sucrose-based ACSF at 34–35°C for 30 min. To minimize sucrose-based depletion of vesicle pools, slices for ultrastructural analysis were additionally transferred to standard ACSF (125 mM NaCl, 25 mM NaHCO_3_, 25 mM D-glucose, 2.5 mM KCl, 1.25 mM NaH_2_PO_4_, 2 mM CaCl_2_, and 1 mM MgCl_2_, equilibrated with 5% CO_2_/95% O_2_, with an osmolarity of ∼316 mOsm) at 34–35°C for at least 5 min before drug application.

### Paired pre-postsynaptic recordings

Slices were placed in a recording chamber and superfused with ACSF for at least 10 min before the onset of recordings. During measurements, slices were superfused with standard ACSF solution. Subcellular patch-clamp recordings from single MFBs and simultaneous recordings from CA3 PNs were performed under an upright microscope with infrared differential interference contrast optics, as previously described (*34*). Pre- and postsynaptic recording pipettes were fabricated from borosilicate glass capillaries of 2.0 mm outer diameter (OD), and 0.6 mm or 1 mm inner diameter (ID) (Hilgenberg, cat#1807528, 1807542), respectively, and had open-tip resistances of 25–35 MΩ and 3–6 MΩ, respectively, when filled with a K^+^-based internal solution (135 mM K-gluconate, 20 mM KCl, 2 mM MgCl_2_, 2 mM Na_2_ATP, 10 mM HEPES, 0.3 mM NaGTP, and 0.1 mM EGTA; for HFS or HFS + LS experiments, modified with 120 mM K-gluconate and 10 mM EGTA in the postsynaptic pipette, pH for both adjusted to 7.28 with KOH, ∼302 mOsm). For selected visualization (Fig. 1A), 0.2% biocytin was added to the presynaptic internal solution, and later developed using streptavidin Alexa Fluor 488 conjugate (Invitrogen, cat#S32354); for the postsynaptic pipette, we included 20 µM biocytin Alexa Fluor 594 (Invitrogen, cat#A12922) to the internal solution. The presynaptic holding potential was set at −70 or −80 mV in the voltage-clamp configuration. APs in MFBs were evoked by brief voltage pulses (amplitude 800–999 mV, duration 0.1–0.2 ms) in the tight-seal, cell-attached configuration. CA3 PNs were held at −70 mV in the whole-cell voltage-clamp configuration for EPSCs recordings. For EPSP recordings, CA3 PNs were injected with hyperpolarizing (Ctrl) or depolarizing (GABA_B_ receptor agonist (R)-baclofen (baclofen, Tocris, cat#0796)) currents and their membrane potentials were held at their average resting membrane potential of approximately −64 mV in the current-clamp configuration (*45*). Membrane potentials of CA3 PNs were closely monitored during EPSP recordings and the holding current was adjusted as required. For experiments with C57BL/6J mice, slices were superfused with standard ACSF followed by ACSF containing 20 µM baclofen for 5 min.

A train stimulation of 15 or 3 stimuli at 50 Hz was delivered every 20 s to the MFB for EPSC or EPSP recordings, respectively. mEPSCs were recorded from CA3 PNs in the presence of 1 µM tetrodotoxin (TTX, Alomone labs, cat#T-550), 10 mM tetraethylammonium (TEA, Sigma-Aldrich, cat#T2265), and 10 µM gabazine (Biotrend, cat#BN0507 and Hello Bio, cat#HB0901). For experiments with Vgat-ires-Cre x Ai32 mice, slices were perfused with ACSF containing 10 µM gabazine for 5 min before the onset of recordings. Slices were stimulated by a train of either 100 light stimuli at 20 Hz or 25 light stimuli at 5 Hz with a CoolLED pE-300^white^ (intensity 5 mW mm^−2^, stimulus duration 5 ms; Filter set 38 HE, Zeiss). For EPSC recordings in initial experiments, a train of 3 electric stimuli was delivered to the MFB either with no light stimulation (Ctrl) or 1 s after 100 light stimuli alternating every 30 s. 5 µM of the GABA_B_ receptor antagonist CGP 55845 hydrochloride (CGP 55845; Hello Bio, cat#HB0960) was added to ACSF to block GABA_B_ receptor activation. Short-term plasticity at MFB–CA3 PNs was induced by delivering a high-frequency stimulation train of 100 electric stimuli at 100 Hz (HFS_100APs_) to MFBs in bouton-attached configuration. A train of 15 electric stimuli was delivered to MFBs every 20 s before and after HFS. For experiments combining HFS and light stimulation, slices were exposed to a train of 100 (at 20 Hz) or 25 (at 5 Hz) light stimuli exactly at 20 s, 40 s, and 60 s after HFS_100APs_, followed by a train of 15 electric stimuli delivered to MFBs 1 s after light stimuli. Postsynaptic series resistance was 9.20 ± 0.58 MΩ (range: 5.2–14.7 MΩ). Series resistance was uncompensated but carefully monitored with a test pulse (5 mV) following each data acquisition sweep. Only recordings with < 300 pA leakage current in the postsynaptic CA3 PN were included in the analyses. Patch-clamp recordings were performed at ∼23.5°C (range: 22–25°C).

### Presynaptic whole-bouton recording

Presynaptic patch pipettes were fabricated from borosilicate glass capillaries (2.0 mm OD, 0.6 mm ID, Hilgenberg, cat#1807528) with an open-tip resistance of 25–30 MΩ, and filled with a Cs^+^-based solution containing 145 mM CsCl, 2 mM MgCl_2_, 2 mM Na_2_ATP, 0.3 mM NaGTP, 5 mM Na_2_-phosphocreatine, 10 mM EGTA, and 10 mM HEPES (pH adjusted to 7.3 with CsOH, osmolarity adjusted to ∼343 mOsm with sucrose). Presynaptic MFBs were voltage-clamped with a holding potential of −80 mV and Ca^2+^ currents were triggered by AP waveforms and recorded, as described in previous publications (*51*, *69*). Ca^2+^ currents were isolated pharmacologically with 1 µM TTX, 20 mM TEA, and 5 mM 4-aminopyridine (4−AP, Sigma-Aldrich, cat#275875) in ACSF. Presynaptic series resistance was 87.0 ± 7.8 MΩ (range: 69.7–119.5 MΩ) before compensation and 40.5 ± 2.2 MΩ (range: 34.1–47.5 MΩ) after compensation (35–65% correction, bandwidth 10 kHz).

### Tissue processing for conventional EM and confocal microscopy

For EM and for confocal microscopy of GABA_B_1 fl/fl x Prox1-Cre mice, two homozygous mice and two wild-type littermate (GABA_B_1 wt/wt x Prox1-Cre) at P21 were anaesthetized with ketamine/xylazine/metamizol (100 mg⋅kg^−1^ ketamine (Ketamidor), VetVita, cat#8-01141; 10 mg⋅kg^−1^ xylazine (Rompun), Elanco Animal Health GmbH, cat#14840; 200 mg⋅kg^−1^ metamizol (Novalgin), Sanofi, cat#3.191; in 0.9% NaCl) and transcardially perfused with 0.1 M phosphate buffer (PB) solution, pH 7.35, followed by ice-cold 4% paraformaldehyde (PFA) in 0.1 M PB. For the EM fixative, 0.05% GA and 15% saturated picric acid solution were added. Brains were dissected, incubated overnight in the fixative solution, and rinsed with 0.1 M PB prior to sectioning. Hippocampal horizontal sections (50–100 μm thickness) were cut in 0.1 M PB using a vibratome (Leica VT1200S).

For EM, tissue sections were washed in PB (3 x 10 min) at RT and incubated in 1% osmium tetroxide in 0.1 M PB for 45 min at RT in the dark. After washing in _dd_H2O (3 x 10 min) at RT, the samples were incubated in 1% uranyl acetate in 50% ethanol at RT for 30 min in the dark. The sections were then dehydrated through a graded ethanol series (50%, 70%, 90%, 96%, 100%, and 100%) for 5–10 min at each concentration, followed by incubation in acetone (2 x 10 min) at RT. For embedding, the samples were sequentially incubated in Durcupan/acetone mixtures at ratios of 1:2 for 1–2 h, 1:1 for 1–2 h, and 2:1 for 1–2 h. After overnight incubation in 100% Durcupan, the samples were embedded and polymerized at 60°C for at least 48 h. Durcupan was prepared by mixing 10.0 g of reagent A, 10.0 g of reagent B, 0.3 g of reagent C, and 0.3 g of reagent D (Sigma-Aldrich, cat#44611, 44612, 44613, and 44614).

For confocal microscopy, sections were washed in 0.1 M PB and blocked and permeabilized with 10% normal goat serum (NGS; Biozol, cat#ENG9010-10) and 0.4% Triton X-100 in 0.1 M PB for 1 h at RT. Then, they were incubated in 5% NGS and 0.4% Triton X-100 in 0.1 M PB with primary antibodies (anti-GABA_B_R_1_, 1:500, guinea pig, gift from A. Kulik; anti-NeuN, 1:500, rabbit, Invitrogen, cat#PA5-37407) overnight at 4°C with gentle shaking. After washing in PB (3 x 10 min), secondary antibodies (goat anti-guinea pig Alexa Fluor 488, 1:500; goat anti-rabbit Alexa Fluor 546, 1:500, both Invitrogen, cat#A-11073 and cat#A-11035) were added in 3% NGS and 0.3% Triton X-100 in 0.1 M PB for 5 h at RT with gentle shaking. Finally, slices were washed again in PB (3 x 10 min) and mounted in Prolong Gold (Invitrogen, cat#P36930). Confocal imaging was performed as described for LICONN iterative expansion microscopy.

### High-pressure freezing (HPF) and freeze substitution (FS) on acute brain slices

For HPF and equivalent to chemical activation of GABA_B_ receptors for patch-clamp recordings, 200 µm thick acute hippocampal slices from C57BL/6J mice were incubated with ACSF containing 20 µM baclofen or with ACSF (Ctrl) for 5 min at 37°C. For flash-and-freeze experiments (*26*), 150-µm-thick acute hippocampal slices from Vgat-ires-Cre x Ai32 mice were incubated with ACSF containing 10 µM gabazine or ACSF for 5 min at 32°C. Following the protocols described in (*26*) and (*35*), slices were loaded into HPF carriers (for drug incubation: type B, 3 mm diameter and 200 μm depth, Wohlwend, cat#662; for flash-and-freeze: type B, 6 mm diameter and 150 μm depth, Wohlwend, cat#1327) enclosed in a droplet of incubation solution. Only instantaneously before freezing, 15% polyvinyl-pyrrolidone in ACSF (PVP, Sigma-Aldrich, cat#PVP10, 37°C, equilibrated with 5% CO_2_/95% O_2_) was used to slightly overfill the carrier. A sapphire disk (3 mm, Wohlwend, cat#405, or 6 mm, Wohlwend, cat#1292) was placed onto the carrier serving as a flat lid and topped by a 300-μm-thick ring (Wohlwend, cat#1241 and cat#1653). The EM ICE (Leica Microsystems) equipped with a blue LED light stimulation module with 460 nm wavelength (LED intensity at specimen 5.5–8.0 mW mm^−2^) was used. For flash-and-freeze experiments, the optical fiber was located at the top of the freezing chamber, thereby illuminating the sample instantaneously before freezing (25 stimuli @ 5 Hz and 100 stimuli @ 20 Hz, respectively, stimulus length 5 ms). All samples were processed at a freezing speed >15000 K s^-1^ and a pressure >2100 bar. FS was conducted in the Leica AFS2 system according to (*35*), with modified incubation time (72 h instead of 96 h) of 0.1% tannic acid (Sigma-Aldrich, cat#403040) and 0.5% GA (Electron Microscopy Sciences, cat#16220) in anhydrous acetone (ChemLab, cat#CL00.2725.1000) embedding in Durcupan resin. This protocol was chosen to enhance vesicle membrane contrast for ET. Samples were positioned flatly in embedding capsule lids and polymerized at 60°C for at least 48 h.

### 2D EM and 3D ET

Sample blocks were trimmed in elongated trapezoids to preserve the entire hippocampal CA3 region (*35*). Both ultrathin sections of 60–100 nm and semi-thin sections of 250 nm were cut with a Histo diamond knife (Diatome AG) at an EM UC7 ultramicrotome (Leica Microsystems). Sections were serially positioned onto Pioloform (5% polyvinyl butyral in trichloromethane, Plano, cat#R1275) coated copper grids (50 mesh and 2×1 mm, Plano, cat#G2050C, G2500C). Positive staining of both section types and carbon coating for semi-thin sections were conducted according to (*35*). 2D electron micrographs from ultra-thin sections were obtained with a standard magnification of 93,000× (pixel size of 0.760 nm) at 80 kV using a transmission electron microscope Thermo Fisher/FEI Tecnai 10 equipped with an OSIS Megaview III G3 camera and using Radius acquisition software.

Tilt image series acquisition was conducted, as described previously (*35*). In brief, image series were acquired at 200 kV using a JEM-2100 (JEOL) electron microscope equipped with a TemCam F416 4k×4k camera (Tietz Video and Image Processing Systems). The acquisition was carried out within a minimum range of −60° to +60° tilt angle with 1° increments at a standardized pixel size of 0.287 nm using the SerialEM software package (RRID:SCR_017293). Tilt-image series alignment and tomographic reconstruction were performed with the ETomo/IMOD software package (RRID:SCR003297) using a patch tracking alignment and weighted back projection algorithm combined with a SIRT-like filter with up to 10^6^ iterations. In selected tomograms with particularly bended presynaptic membranes, a Wiener-like deconvolution filter was additionally applied after reconstruction using ETomo. Segmentation and 3D reconstruction of electron tomograms were carried out in the 3Dmod/IMOD software package (RRID:SCR003297).

### Sample selection for 2D EM and 3D ET

Samples were selected from experiments with interleaved design. Both imaging modalities (2D, 3D) were applied to the same sample. Prior to selection, samples were systematically examined for potential freezing and embedding artefacts using ultra-thin sections according to (*26*, *35*). The selection for image acquisition was equivalent to (*26*, *35*). In brief, we acquired electron micrographs and tilt series of AZs in 2–6 µm giant MFBs, preferably in the CA3b *stratum lucidum*. Only boutons containing clearly identifiable CA3b pyramidal spine heads, some with a spine apparatus, and postsynaptic densities (PSDs) directly opposite to AZs were analyzed. AZs were defined by presynaptic clusters of docked synaptic vesicles opposed to postsynaptic membranes, separated by the synaptic cleft. Non-MFB–CA3b spine contacts were excluded. Each sample was identified by a freeze substitution ID blinded to its experimental condition. All electron micrographs and reconstructed tomograms were randomized by random number sampling prior to docked vesicle quantification and segmentation, respectively.

### LICONN iterative expansion microscopy

Transcardial fixative perfusion, brain tissue processing, epoxide treatment, iterative polymerization of hydrogels, immunolabeling, and spinning-disc confocal microscopy were performed according to (*60*). Briefly, three C57BL/6J mice (P21) were transcardially perfused with 4% paraformaldehyde (TAAB, cat#P001-1) and 10% acrylamide (Sigma-Aldrich, cat#A9099) in phosphate-buffered solution (PBS). For further processing, 50-μm brain slices were cut based on previously described criteria. Before polymerization of the first expandable hydrogel, conducted according to (*60*), brain slices were trimmed to the region of the hippocampus. After gelation for 2 h at 37°C, hydrogel-tissue hybrids were trimmed to a region of interest containing only the dentate gyrus and CA3, and denatured for 100 min at 95°C. After neutralization of unreacted groups from the first gelation (*60*), hydrogels underwent immunolabeling with the following specifications: 1. Primary antibodies: anti-Munc13-1, 1:300, rabbit (Synaptic Systems, cat#126103); anti-VGAT cytoplasmic domain, 1:300, guinea pig (Synaptic Systems, cat#131004), in PBS at 4 °C overnight with gentle agitation. 2. Secondary antibodies: goat anti-rabbit Alexa Fluor 546, 1:300 (Invitrogen, cat#A-11035); goat anti-guinea pig STARRED, 1:150 (Abberior, cat#stred-1006), in PBS at 4°C overnight with gentle agitation. Evaluation of immunolabeling was performed using hydrogels re-expanded in milli-Q water. Further sample processing including stabilizing hydrogel polymerization, neutralization, and second expandable hydrogel polymerization was conducted according to (*60*). Before imaging, pan-protein staining was performed using overnight incubation of 40 μM NHS-Atto 488 (Atto-Tec, cat#AD-488-31) in PBS at 4°C under gentle agitation. Confocal imaging was performed using an Andor Dragonfly spinning-disc microscope equipped with a Nikon Ti2E inverted stand, a motorized stage, an Andor Zyla 4.2-megapixel sCMOS camera (2048×2048 px), and a 40× water-immersion objective (Nikon Apochromat LWD 40× lambda S, NA 1.15, water, working distance 0.6 mm). Image volumes were acquired using Andor Fusion software v2.2. Color channels with excitation wavelengths of 488 nm (pan-protein), 561 nm (Munc13-1), and 637 nm (VGAT) were recorded sequentially at each Z-plane. All image volumes were obtained in hippocampal CA3 *stratum lucidum* and within one imaging session using hydrogels prepared in one batch.

### Electrophysiology data analysis

Patch-clamp data were acquired with a MultiClamp 700B amplifier (Molecular Devices) and MultiClamp 700B Commander (RRID:SCR_018455) and digitized using a CED Power1401 interface (Cambridge Electronic Design). Signals were low-pass filtered at 10 kHz (Bessel) and sampled at 50 kHz. Pulse generation and data acquisition were performed using Igor Pro v6.37 (WaveMetrics, RRID:SCR_000325) and FPulse version 3.45 (U. Fröbe, University of Freiburg, RRID:SCR_027414) with custom FPulse scripts. EPSC and EPSP amplitudes and kinetics were analyzed using Stimfit version 0.15.8 (RRID:SCR_016050) or custom Matlab scripts (Matlab R2022b, MathWorks, RRID:SCR_001622). For EPSC and EPSP peak detection, the analysis window was set between 1 and 10 ms after the stimulus. Pool analysis was performed by linear regression of cumulative EPSC amplitudes (SMN method) or fitting a product of an exponential facilitation and an exponential depression function (TR method, with custom Mathematica script (Mathematica v14.0, Wolfram Research, RRID:SCR_014448), as previously described (*4*, *46*). For each experiment involving HFS, the trace corresponding to the time point (20 s, 40 s, or 60 s after HFS) with the largest EPSC_1_ peak amplitude was used for analyses of EPSC amplitude and pool parameters during short-term plasticity. mESPC events were detected using a template-matching method (*87*), including the evaluation of detection limits (*88*). Ca^2+^ current recordings were filtered with a 5-kHz low-pass digital filter (Gaussian). Capacitive and leakage currents from Ca^2+^ current recordings were subtracted using the pipette capacitance compensation circuit of the amplifier and a P over −4 correction method. EPSC and Ca^2+^ current amplitudes in Ctrl condition were tested for stationarity and only recordings without a statistically significant rundown (Spearman rank correlation coefficient with *p* > 0.05) were included for further analysis.

### Ultrastructural data analysis

AZs were defined as presynaptic compartments containing dense vesicle clusters including docked vesicles in near proximity to and at the presynaptic membrane. The presynaptic 2D AZ profile and 3D AZ surface, respectively, were defined by their localization opposite to the postsynaptic membrane at which electron-dense postsynaptic filaments were densely connected. Pre- and postsynaptic compartments were separated by the synaptic cleft, a characteristic widening between both membranes containing a network of electron-dense filaments. Synaptic vesicles situated within 0 to 2-nm distance to the presynaptic membrane and in direct connection with the presynaptic membrane were defined as ‘docked’ (*53*). To align with previous docking criteria in 2D AZ profile analysis (0–5 nm; (*26*)), we additionally quantified docking within this extended range.

For 2D transmission EM, quantification of vesicle radii and distances to presynaptic membranes was carried out using Fiji Version v1.54p (RRID: SCR_002285). For 3D ET, annotation of objects and export of both 3D coordinates and further geometrical parameters were conducted according to (*35*). In brief, vesicles of 0–50 nm Euclidean distance to the presynaptic membrane were included into the annotation (*26*, *35*). Presynaptic membranes (AZ profiles) were two-dimensionally annotated as ‘open’ line objects with linear interpolation every 15 virtual sections. 3D coordinates of synaptic vesicle centers and membranes, vesicle radii, contour lengths, object areas, and volumes were extracted from IMOD models. The data were processed in Jupyter Notebook (v8.6.2, RRID: SCR_018315) using Python (v3.13.0, RRID:SCR_008394) with custom-written scripts based on previously published code (*35*). For the Monte Carlo envelope (Fig. 5I and J), observed NNDs (Fig. 5G) were defined within a 2D rectangle with AZ-like dimensions, consisting of the mean AZ profile length in both conditions and an electron tomographic section thickness of 250 nm. A thickness of 200 nm was also tested to account for beam-induced section shrinkage, yielding similar results. For both conditions, the median vesicle density (synaptic vesicles per 0.01 μm² AZ area) was used to determine the number of points for null model simulations. In the MC simulation, the null model consisted of M = 999 independent simulations of random point patterns within the rectangular region. For each simulation, the specified number of points, defined by the mean density and AZ area, was randomly distributed using uniform sampling. NNDs were computed for each simulated point pattern using KDTree from SciPy (v1.16.1, RRID:SCR_008058). The empirical cumulative distribution function (CDF) of all NNDs was computed, compared to and plotted against the Monte Carlo envelope to visualize deviations from complete spatial randomness, based on (*89*). A 95% confidence envelope was constructed from the simulated CDFs using percentiles (i) α/2, (ii) 1-α/2, with α = 0.05). Random number generation was seeded for reproducibility.

### Expansion microscopy data analysis

First, raw image volumes underwent denoising using Noise2Void (v0.3.2, RRID:SCR_027415). For each image volume, 10,000 patches with a size of 64^3^ pixels were randomly sampled without augmentation. We trained for 60 epochs with 500 training steps per epoch and a batch size of 16 patches. For manual segmentation in Paintera (v1.10.2, RRID:SCR_027413), five MFBs per image volume (n = 6 volumes) were selected based on the morphological criteria specified for the EM analysis. Mean dimensions of the image volumes were as follows: x = 20.30 µm; y = 20.30 µm; z = 10.66 ± 0.39 µm/10.48 (9.59–11.86) µm. The absence of pan-protein labeling within the MFB, consistent with the dense synaptic vesicle pool, served as an additional identification criterion. Segmentation was cross-referenced against presynaptic AZ Munc13-1 puncta in the raw image volumes using Imaris (v9.9.1, Oxford instruments, RRID:SCR_007370). VGAT cluster identification and distance measurement were performed using a custom written Python (v3.11) script. For this, the VGAT channel was imported from each image volume with the following px sizes: x = 0.305 µm, y = 0.305 µm, z = 0.798 µm. Prior to peak detection, a Difference of Gaussians (DoG) filter was applied to enhance fluorescent puncta of ∼3 pixel diameter using σ_1_ = (1.24, 1.24, 0.48) px and σ_2_ = 1.6 × σ_1_ px (x, y, z). The resulting DoG filtered volumes were normalized, setting their minimum intensity value to 0 and the 99.9^th^ percentile to 1. Then, local maxima of fluorescent puncta were detected with a threshold of 0.5 and minimum distance of 1 px. Detections at the volume borders were excluded. Detected peaks were clustered using the Python implementation (skicit-learn, RRID:SCR_002577) of density-based spatial clustering of applications with noise (DBSCAN; eps = 3.5, min_samples = 4). DBSCAN-detected VGAT clusters (mean 699.83 ± 65.10 / median 663.00 (489.00–916.00)) were manually proofread to generate a sample set of semi-automatically detected clusters (mean 280.67 ± 21.82 / median 253.50 (236.00–365.00)) using a custom extension on napari (v0.6.2, RRID:SCR_022765) fulfilling both of the following criteria: i) tissue context (interneuron terminal-like structures) provided by pan-protein staining, and ii) the co-expression of Munc13-1 puncta. An expansion factor of 15.44 was used to calculate pre-expansion distances (*60*). All values reported in Fig. 6 and Fig. S7 are pre-expansion. NNDs between VGAT clusters and individual segmented MFBs were calculated as the NND between the 3D center of mass (COM) of each cluster and the nearest surface of the MFB segmentation (MFB outer membrane).

### Statistical analysis

Statistical analysis was carried out with the statistical software programs Sigma Plot 14 (Systat Software, RRID:SCR_003210), OriginPro 2025 (Origin Lab, RRID:SCR_014212), RStudio v2025.05.1 (Posit Software, RRID:SCR_001905), and GraphPad Prism v10.2.2 (GraphPad Software, RRID:SCR_000306). For all data, normality was tested using Shapiro-Wilk tests and Kolmogorov-Smirnov-tests. For comparison of two groups, two-sided paired non-parametric Wilcoxon signed-rank, aligned rank transform for non-parametric factorial ANOVA with contrast (Holm adjustment based on the low sample number), two-sided unpaired non-parametric Mann-Whitney rank sum, and parametric student’s t-test were used. Power analyses were performed separately and considered in the interpretation as appropriate. For maximum likelihood fitting, the frequency distribution of mEPSC 20–80% rise time was fitted with a two-component Gaussian model on unbinned data. For testing stationarity and data correlation, a Spearman rank correlation coefficient was calculated. The experimental procedure was independently replicated several times within the laboratory. n/N was used as an abbreviation for the sample number. Data are reported as mean ± SEM (electrophysiology/expansion microscopy) and median with 10^th^/90^th^ percentiles (EM/ET), unless otherwise indicated. Respectively, box plots depict data as median (solid line), mean (dashed line), 25^th^/75^th^ quartile (box), and for EM data 10^th^/90^th^ percentile (whiskers), as well as min/max points (whiskers) for electrophysiology. Scatter plots show individual data points unless indicated otherwise. All graphs were produced with OriginPro 2025 and GraphPad Prism. Figures were assembled using Affinity Designer 2 v2.6.3 (Serif, RRID:SCR_016952). Tables S1–6 contain all numerical values not stated in text and figure legends.

## Acknowledgments

This research was supported by the Scientific Services Units of ISTA (Electron Microscopy Facility, Imaging and Optics Facility, Scientific Computing, Preclinical Facility, and Miba Machine Shop) and the Imaging Core Facility of the University of Würzburg, in particular by microscope access. The JEOL JEM-2100 transmission electron microscope used in this study was funded by the Deutsche Forschungsgemeinschaft (DFG, German Research Foundation) – 218894163. We thank C. Stigloher and T. Vogels for critically reading the manuscript. We are grateful to F. Marr, C. Altmutter, M. Ederer, and N. Valente for excellent technical assistance, and E. Kralli-Beller for manuscript editing. For data analysis support, we are grateful to A. Schlögl (electrophysiology) and C. Sommer (expansion microscopy). For ultramicrotomy support, we are grateful to V. Zheden. We thank B. Bettler for the GABA_B_1 floxed mice and A. Kulik for the GABA_B_1 antibody. Part of the work has been published in P.L.’s thesis in partial fulfillment of the requirements for the degree of Doctor of Philosophy.

## Funding

ERC Advanced Grant 692692 “GIANTSYN” (P.J.)

ERC Advanced Grant 101199096 “CA3-SYNGRAM” (P.J.)

Fonds zur Förderung der wissenschaftlichen Forschung P 36232-B, stand-alone grant (P.J.)

Fonds zur Förderung der wissenschaftlichen Forschung PAT 4178023, principal investigator project (P.J.)

Fonds zur Förderung der wissenschaftlichen Forschung 10.55776/CoE 16 “GABA neurons” (P.J.)

European Union’s Horizon 2020 research and innovation programme under the Marie Skłodowska-Curie grant agreement No 101034413 (K.L.)

## Author contributions

Conceptualization: P.J., K.L., Y.O.

Methodology: all authors

Investigation: P.L. (electrophysiology; expansion microscopy), K.L. (electron microscopy/tomography), M.R.T. (contribution to expansion microscopy), J.G.D. (contribution to expansion microscopy)

Formal analysis: P.L., K.L., P.J.

Visualization: P.L., K.L.

Writing—original draft: K.L., P.J.

Writing—review & editing: all authors

Funding acquisition: P.J., K.L.

Supervision: P.J.

## Data, code, and materials availability

All data and code needed to evaluate and reproduce the results in the paper are present in the paper and/or the Supplementary Materials. This study did not generate any new materials.

