## Supplemental Material for "Presynaptic inhibition switches hippocampal mossy fiber computations and limits vesicle pool plasticity"

Peipeng Lin *et al.*

**This PDF file includes:**

Figs. S1 to S8
Tables S1 to S7

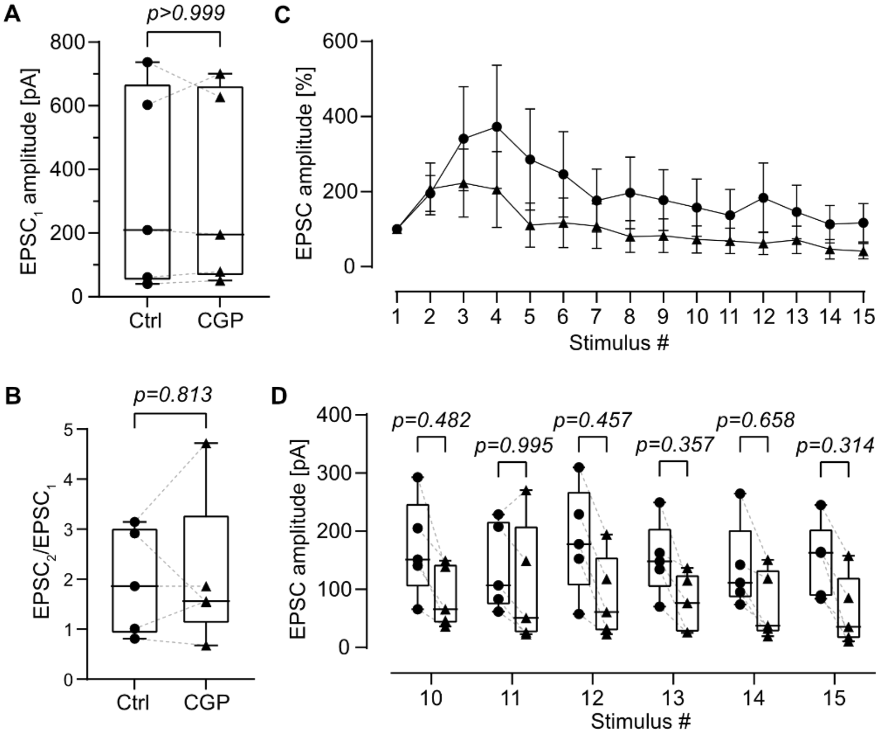

**Fig. S1, related to Fig. 1. Effects of 5 µM of the $GABA_B$ receptor antagonist CGP 55845 on MFB–CA3 PN synaptic transmission. (A, B)** Summary box plots of $EPSC_1$ peak amplitude and paired-pulse ratio ($EPSC_2/EPSC_1$). CGP 55845 has no detectable effect on both parameters, suggesting a lack of tonic activation of presynaptic $GABA_B$ receptors and constitutive GIRK channel activity. **(C, D)** Normalized and absolute EPSC peak amplitudes plotted against stimulus number. CGP 55845 has no detectable effect on synaptic dynamics or EPSC peak amplitudes at late stimuli of a 50-Hz train, suggesting that $GABA_B$ receptors are not recruited via an MFB → interneuron → CA3 PN disynaptic inhibition pathway. N = 5 MFB–CA3 PN pairs.

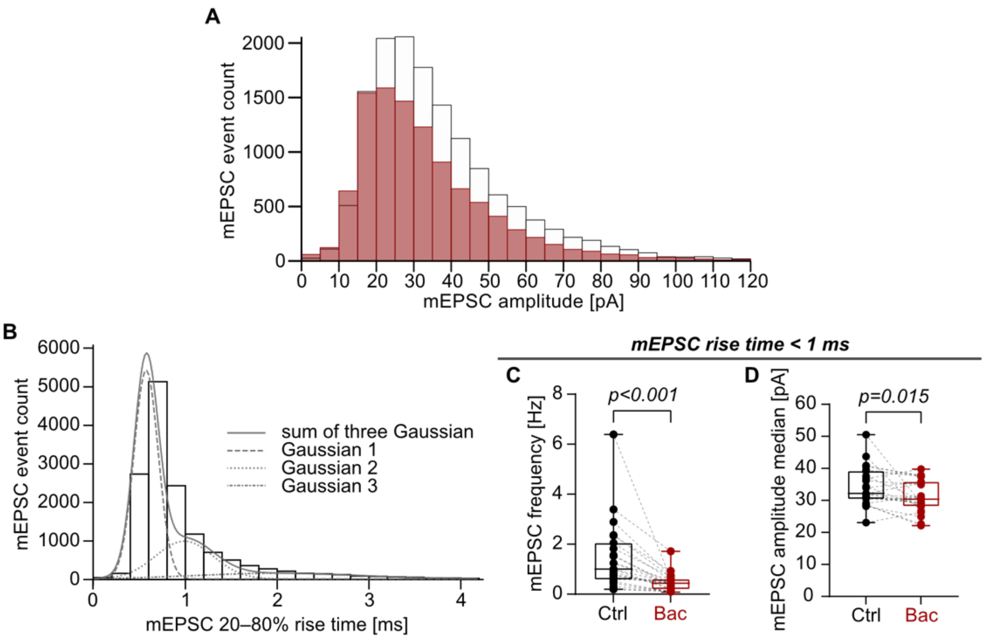

**Fig. S2, related to Fig. 1. Effects of 20 µM baclofen on mEPSCs at MFB–CA3 PN synapses. (A)** Frequency distribution of mEPSC amplitudes in Ctrl and Bac. Bin width = 5 pA. **(B)** Frequency distribution of mEPSC 20–80% rise time analyzed with a three-component Gaussian model using maximum likelihood fitting of unbinned data, suggesting that mEPSC events with fast kinetics (20–80% rise time < 1 ms) are likely of mossy fiber origin. Bin width = 0.2 ms. **(C, D)** Summary box plots of mEPSC frequency and mEPSC amplitude median only including events with 20–80% rise time < 1 ms in Ctrl and Bac, suggesting a presynaptic effect of $GABA_B$ receptor activation at MFB–CA3 PN synapses. N = 23 CA3 PNs.

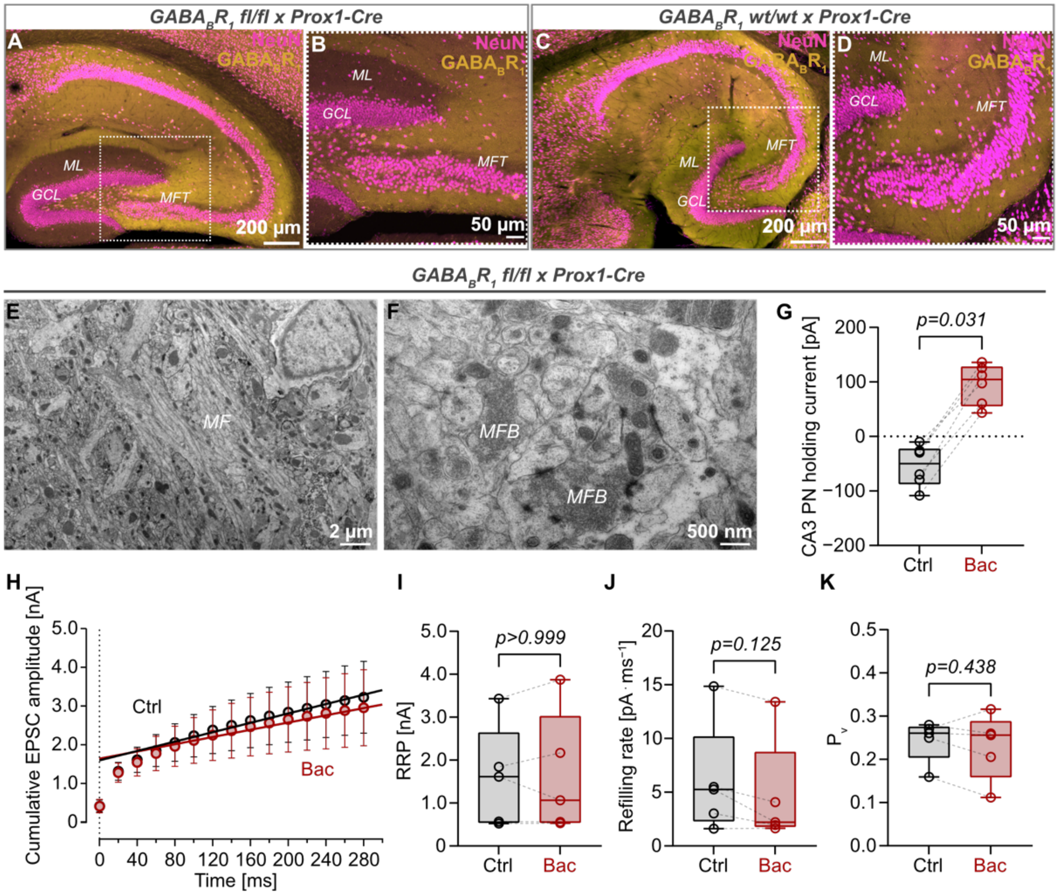

**Fig. S3, related to Fig. 1. Effects of 20 µM baclofen on mossy fiber synaptic transmission in GABA$_B$R$_1$ fl/fl x Prox1-Cre mice. (A–D)** Maximum intensity Z-projection of confocal images of the hippocampal formation from a homozygous GABA$_B$R$_1$ fl/fl x Prox1-Cre mouse **(A, B)** and its wild-type littermate **(C, D)** stained for neuronal nuclei (NeuN, magenta) and the GABA$_B$R$_1$ (yellow). Images from the conditional knock-out mouse **(A, B)** present with diminished GABA$_B$R$_1$ signal in the molecular layer and granule cell layer in dentate gyrus as well as alongside the mossy fiber tract compared to controls **(C, D)**. **(E–F)** Representative 2D electron micrographs of 70-nm ultrathin, classically fixed tissue sections of the ultrastructurally intact mossy fiber tract with MFBs in CA3 of a GABA$_B$R$_1$ fl/fl x Prox1-Cre mouse. **(G)** Summary box plot of the holding current of CA3 PN in Ctrl and in Bac. **(H)** Cumulative plot of EPSC peak amplitudes evoked by a 50-Hz train of 15 stimuli in Ctrl and Bac. RRP size, refilling rate, and $P_v$ were estimated by linear regression of the last four data points and back-extrapolation to time 0. **(I–K)** Summary box plots of RRP size, refilling rate, and $P_v$, indicating no effect of baclofen and, thus, the specificity of the observed GABA$_B$R-mediated presynaptic inhibition. Data points from the same MFB–CA3 PN pair are connected by dashed lines. Data in G from N = 6, data in H–K from N = 5 MFB–CA3 PN pairs.

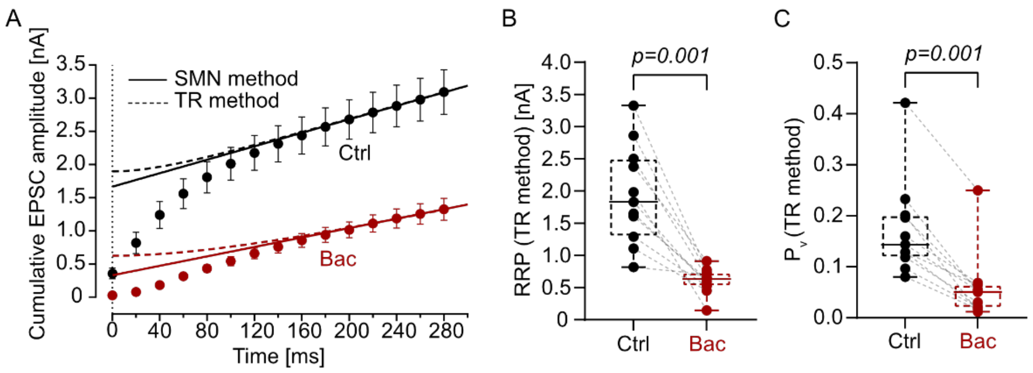

**Fig. S4, related to Fig 3. Analysis of RRP size and $P_v$ using a method accounting for a time-dependent refilling rate. (A)** Cumulative plot of EPSC peak amplitudes evoked by a 50-Hz train of 15 stimuli in Ctrl and Bac. Solid lines show linear fit according to the SMN method (*46*), assuming a constant refilling rate. Dashed lines indicate fit according to the TR method (*47*), assuming that refilling rate increases as release sites are vacated. **(B, C)** Summary box plots of RRP size and $P_v$ in Ctrl and Bac according to the TR method, showing significant reduction in both parameters.

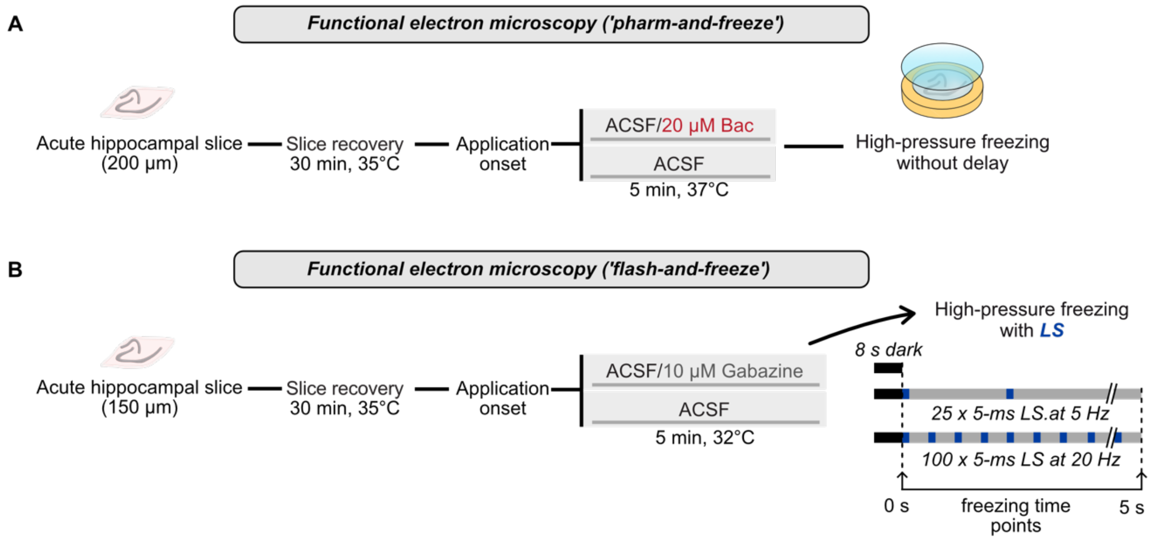

**Fig. S5, related to Fig. 4, Fig. 5, and Fig. 8. Experimental workflow for functional electron microscopy. (A)** Scheme of the high-pressure freezing experiment for $GABA_B$ receptor activation in MFBs using baclofen ('pharm-and-freeze'). **(B)** Scheme of the high-pressure freezing experiment for $GABA_B$ receptor activation in MFBs using light stimulation of VGAT-positive interneurons ('flash-and-freeze').

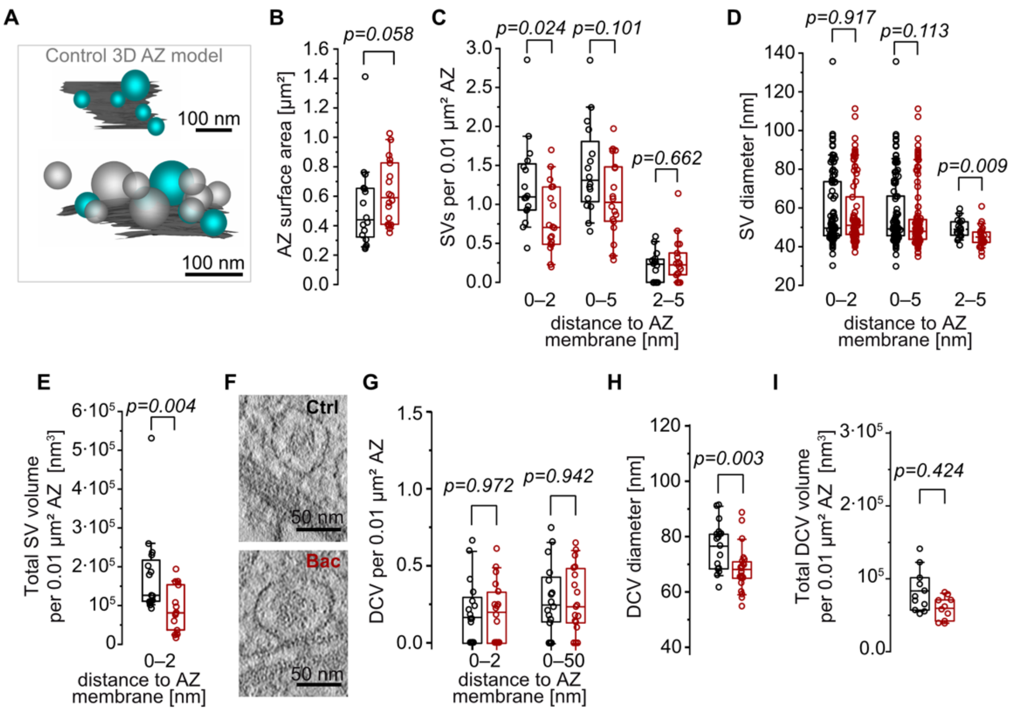

**Fig. S6, related to Fig. 5. Ultrastructural characterization of 3D vesicle pools at MFB AZs using ET. (A)** Representative 3D model of a Ctrl AZ (dark gray) presented in Fig. 5B, containing total SV pool (0–50 nm; membrane-proximal pool; light gray) and tightly docked clear SVs (0–2 nm; turquoise). The inset depicts the orthogonal view. Further analysis of ultrastructural features at MFB AZs supported a specific effect of Bac on the tightly docked vesicle pool. AZ surface area **(B)** and docked clear SV number per 0.01 µm² AZ surface area **(C)**, using a broader docking definition (0–5 nm) and further differentiation in docking subclasses ('loosely docked', 2–5 nm), remained largely unaffected compared to Ctrl. For comparison, the plot of the number of docked clear vesicles (0–2 nm) per 0.01 µm² AZ surface area from Fig. 5F is repeated in the summary box plot. **(D)** Summary box plots of 2D docked clear SV diameter obtained from ET integrating the docking definition from panel **(C)**, revealing a slight diameter effect on the small fraction of loosely docked vesicles. **(E)** Summary box plot of the total docked clear SV volume normalized to the AZ surface area in Ctrl and Bac. **(F)** Representative electron tomographic slices highlighting dense-core vesicles (DCVs) in Ctrl (top) and Bac (bottom). **(G)** Summary box plots of the number of docked DCVs and total DCV number (0–50 nm distance to the AZ membrane) per 0.01 µm² AZ surface area, revealing no reduction under baclofen. **(H, I)** Summary box plots of 2D ET docked DCV diameter and total docked DCV volume normalized to the AZ surface area in Ctrl and Bac, collectively revealing no major effect of baclofen on the dense core vesicle pool.

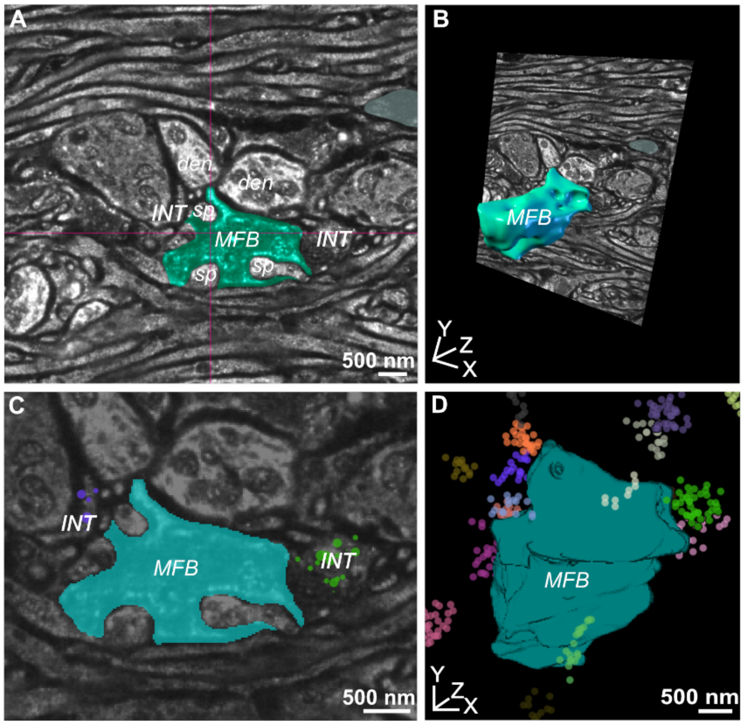

**Fig. S7, related to Fig. 6. Representative analysis of a single image volume obtained with ~16-fold LICONN expansion microscopy. (A, B)** Manual segmentation of the same MFB in **Fig. 6** based on tissue context revealed by pan-protein staining in Paintera (**A**) and a 3D rendering of the segmentation (**B**). Scale bars reflect pre-expansion lengths. Abbreviations: INT, interneuron terminal; den, dendrite; sp, spine head. **(C, D)** Semi-automatically detected VGAT clusters identified in close proximity to the annotated MFB in a single 2D plane (**C**) and in 3D (**D**) using napari.

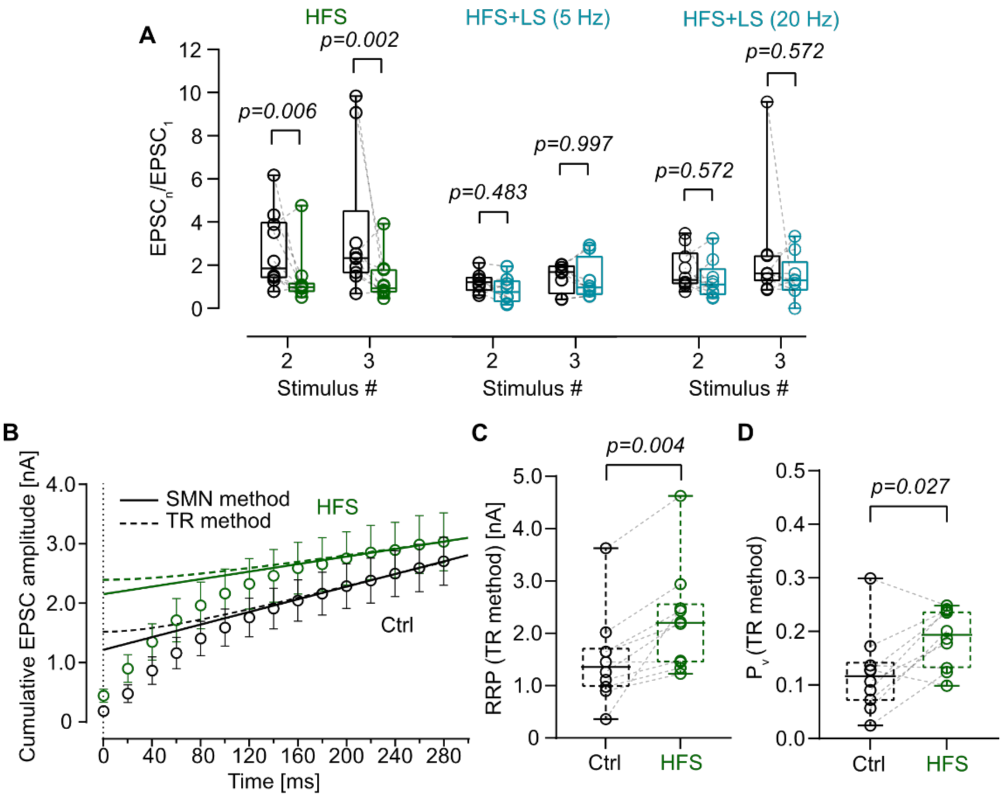

**Fig. S8, related to Fig. 9 and Fig. 10. Analysis of HFS-induced changes in synaptic dynamics and vesicle pool parameters using the TR method. (A)** Summary box plot for $EPSC_n/EPSC_1$ for stimulus number 2 and 3 in Ctrl versus HFS, Ctrl versus HFS+LS (5 Hz), and Ctrl versus HFS+LS (20 Hz). N = 10 (HFS), 8 (HFS+LS (5 Hz)), and 10 (HFS+LS (20 Hz)) MFB–CA3 PN pairs. **(B)** Cumulative plot of EPSC peak amplitudes evoked by a 50-Hz train of 15 stimuli in Ctrl and HFS. The last 4 data points were fitted with SMN (solid lines) and TR methods (dashed lines) to estimate RRP size and $P_v$. **(C, D)** Summary box plots of RRP size (**C**) and $P_v$ (**D**), in Ctrl and HFS estimated using the TR method, suggesting an expansion of the presynaptic vesicle pool and an increase in release probability. N = 10 MFB-CA3 PN pairs.

**Table S1.**

| Parameter [unit] | Mean ± SEM / median ($25^{th}$–$75^{th}$ percentile) | p-value and statistical test | Fig. |
|---|---|---|---|
| $EPSC_1$ peak amplitude [pA] | Ctrl: 358.29 ± 80.61 / 235.34 (159.34–575.98) | *p* = 0.001; Wilcoxon signed rank test | 1D |
| | Bac: 27.13 ± 4.88 / 27.97 (10.76–41.05) | | |
| $EPSC_1$ failure proportion | Ctrl: 1.82 ± 1.22 / 0 (0–0) | *p* = 0.001; Wilcoxon signed rank test | 1E |
| | Bac: 48.79 ± 7.15 / 50 (26.67–70) | | |
| $EPSC_1$ latency [ms] | Ctrl: 1.44 ± 0.05 / 1.53 (1.26–1.56) | *p* = 0.001; Wilcoxon signed rank test | 1F |
| | Bac: 1.95 ± 0.10 / 1.90 (1.60–2.13) | | |
| $EPSC_1$ 20–80% rise time [ms] | Ctrl: 1.07 ± 0.08 / 0.95 (0.83–1.25) | *p* = 0.005; Wilcoxon signed rank test | 1G |
| | Bac: 0.68 ± 0.13 / 0.58 (0.39–0.96) | | |
| $EPSC_1$ decay **τ** [ms] | Ctrl: 5.74 ± 0.24 / 5.37 (5.19–6.57) | *p* = 0.007; Wilcoxon signed rank test | 1H |
| | Bac: 4.20 ± 0.44 / 4.78 (2.58–5.27) | | |
| Norm. $CV^{-2}$ vs Norm. $EPSC_1$ amplitude | Norm. $CV^{-2}$: 0.10 ± 0.02 | N/A | 1I |
| | Norm. $EPSC_1$ amplitude: 0.08 ± 0.02 | | |

| | | | |
|---|---|---|---|
| $EPSC_2$ / $EPSC_1$ | Ctrl: 1.55 ± 0.19 / 1.42 (1.24–1.80) | $p = 0.459$; Aligned rank transform for non-parametric factorial (ART) ANOVA with contrast test (Holm adjustment) | 1J |
| | Bac: 2.40 ± 0.47 / 2.04 (1.34–2.41) | | |
| $EPSC_3$ / $EPSC_1$ | Ctrl: 1.63 ± 0.27 / 1.28 (1.05–2.33) | $p < 0.0001$; ART ANOVA with contrast test (Holm adjustment) | |
| | Bac: 4.79 ± 0.78 / 4.76 (2.85–6.53) | | |
| RRP [nA] | Ctrl: 1.70 ± 0.23 / 1.43 (1.11–2.25) | $p = 0.001$; Wilcoxon signed rank test | 3B |
| | Bac: 0.33 ± 0.06 / 0.29 (0.18–0.48) | | |
| Refilling rate [pA $ms^{-1}$] | Ctrl: 5.18 ± 0.72 / 4.20 (3.45–7.53) | $p = 0.005$; Wilcoxon signed rank test | 3C |
| | Bac: 3.55 ± 0.63 / 2.90 (2.30–4.68) | | |
| $P_v$ | Ctrl: 0.20 ± 0.03 / 0.18 (0.14–0.22) | $p = 0.002$; Wilcoxon signed rank test | 3D |
| | Bac: 0.10 ± 0.02 / 0.08 (0.04–0.14) | | |
| RRP (TR method) [nA] | Ctrl: 1.94 ± 0.23 / 1.83 (1.29–2.51) | $p = 0.001$; Wilcoxon signed rank test | S4B |
| | Bac: 0.61 ± 0.06 / 0.64 (0.52–0.73) | | |
| $P_v$ (TR method) | Ctrl: 0.17 ± 0.03 / 0.14 (0.12–0.20) | $p = 0.001$; Wilcoxon signed rank test | S4C |
| | Bac: 0.06 ± 0.02 / 0.05 (0.02–0.06) | | |
| $Ca^{2+}$ current peak amplitude [pA] | Ctrl: 115.27 ± 20.91 / 100.66 (72.70–164.03) | $p = 0.031$; Wilcoxon signed rank test | 3G |

| | | | |
|---|---|---|---|
| | Bac: 75.95 ± 14.43 / 73.08 (41.41–112.18) | | |
| $Ca^{2+}$ total charge [fC] | Ctrl: 78.53 ± 12.95 / 65.22 (53.87–111.94) | *p* = 0.031; Wilcoxon signed rank test | 3H |
| | Bac: 46.97 ± 7.71 / 45.21 (28.98–64.94) | | |
| $EPSC_1$ peak amplitude [pA] (CGP) | Ctrl: 330.09 ± 143.29 / 209.70 (50.61–669.77) | $p > 0.999$; Wilcoxon signed rank test | S1A |
| | Bac: 330.78 ± 138.75 / 195.34 (65.04–664.24) | | |
| $EPSC_2$ / $EPSC_1$ (CGP) | Ctrl: 1.95 ± 0.48 / 1.86 (0.91–3.03) | *p* = 0.813; Wilcoxon signed rank test | S1B |
| | Bac: 2.07 ± 0.69 / 1.56 (1.11–3.29) | | |
| $EPSC_1$ peak amplitude [pA] ($GABA_BR1$ fl/fl x Prox1-Cre) | Ctrl: 361.59 ± 140.45 / 289.22 (79.06–600.49) | $p > 0.999$; Wilcoxon signed rank test | 1N |
| | Bac: 357.89 ± 151.47 / 225.47 (70.97–668.55) | | |
| CA3 PN resting potential [mV] ($GABA_BR1$ fl/fl x Prox1-Cre) | Ctrl: -62.12 ± 2.13 / -64.40 (-66.20—56.525) | *p* = 0.031; Wilcoxon signed rank test | 1O |
| | CGP: -80.42 ± 2.19 / 79.95 (-85.03—77.38) | | |

| | | | |
|---|---|---|---|
| CA3 PN holding current [pA] ($GABA_BR1$ fl/fl x Prox1-Cre) | Ctrl: -54.25 ± 15.41 / -50.00 (-87.10– -22.78) | *p* = 0.031; Wilcoxon signed rank test | S3G |
| | Bac: 95.50 ± 15.02 / 104.55 (55.78–127.70) | | |
| RRP [nA] ($GABA_BR1$ fl/fl x Prox1-Cre) | Ctrl: 1.60 ± 0.53 / 1.61 (0.54–2.64) | *p* > 0.999; Wilcoxon signed rank test | S3I |
| | Bac: 1.64 ± 0.63 / 1.06 (0.54–3.02) | | |
| Refilling rate [pA $ms^{-1}$] ($GABA_BR1$ fl/fl x Prox1-Cre) | Ctrl: 6.05 ± 2.32 / 5.26 (2.31–10.17) | *p* = 0.125; Wilcoxon signed rank test | S3J |
| | Bac: 4.66 ± 2.23 / 2.21 (1.79–8.74) | | |
| $P_v$ ($GABA_BR1$ fl/fl x Prox1-Cre) | Ctrl: 0.24 ± 0.02 / 0.26 (0.20–0.28) | *p* = 0.438; Wilcoxon signed rank test | S3K |
| | Bac: 0.23 ± 0.03 / 0.26 (0.16–0.29) | | |
| $EPSC_1$ peak amplitude [pA] (LS (5 Hz)) | Non-LS: 410.62 ± 82.69 / 304.27 (215.78–637.11) | *p* = 0.002; Wilcoxon signed rank test | 7J |
| | LS (5 Hz): 282.66 ± 65.47 / 224.37 (119.39–484.24) | | |
| $EPSC_1$ peak amplitude [pA] (LS (20 Hz)) | Non-LS: 368.06 ± 97.30 / 256.17 (193.05–577.34) | *p* = 0.008; Wilcoxon signed rank test | |
| | LS (20 Hz): 213.38 ± 91.80 / 112.43 (89.52–228.33) | | |

| | | | |
|---|---|---|---|
| $EPSC_1$ peak amplitude [pA] (LS (CGP)) | Non-LS (CGP): 426.87 ± 126.43 / 373.85 (165.77–883.43) | $p = 0.938$; Wilcoxon signed rank test | |
| | LS (CGP): 416.30 ± 133.16 / 300.88 (182.40–885.94) | | |
| $EPSC_2$ / $EPSC_1$ (LS (5 Hz)) | Non-LS: 1.25 ± 0.19 / 1.23 (0.76–1.46) | $p = 0.317$; ART ANOVA with contrast test (Holm adjustment) | 7K |
| | LS (5 Hz): 1.70 ± 0.28 / 1.48 (0.90–2.47) | | |
| $EPSC_3$ / $EPSC_1$ (LS (5 Hz)) | Non-LS: 1.78 ± 0.36 / 1.59 (0.98–2.55) | $p = 0.038$; ART ANOVA with contrast test (Holm adjustment) | |
| | LS (5 Hz): 3.27 ± 0.79 / 2.71 (2.02–4.01) | | |
| $EPSC_2$ / $EPSC_1$ (LS (20 Hz)) | Non-LS: 1.72 ± 0.28 / 1.28 (1.10–2.44) | $p = 0.130$; ART ANOVA with contrast test (Holm adjustment) | |
| | LS (20 Hz): 2.82 ± 0.54 / 2.88 (1.40–3.49) | | |
| $EPSC_3$ / $EPSC_1$ (LS (20 Hz)) | Non-LS: 1.78 ± 0.36 / 1.59 (0.98–2.55) | $p = 0.121$; ART ANOVA with contrast test (Holm adjustment) | |
| | LS (20 Hz): 3.27 ± 0.79 / 2.71 (2.02–4.01) | | |
| $EPSC_2$ / $EPSC_1$ (LS (CGP)) | Non-LS (CGP): 1.91 ± 0.32 / 2.07 (1.14–2.84) | $p > 0.999$; ART ANOVA with contrast test (Holm adjustment) | |

| | | |
|---|---|---|
| | LS (CGP): 1.92 ± 0.44 / 1.90 (0.82–3.36) | |
| $EPSC_3$ / $EPSC_1$ (LS (CGP)) | Non-LS (CGP): 1.69 ± 0.65 / 1.08 (0.45–2.01) | $p > 0.999$; ART ANOVA with contrast test (Holm adjustment) |
| | LS (CGP): 1.65 ± 0.74 / 1.23 (0.37–1.56) | |

Effects of 20 μM baclofen and synaptically released GABA on evoked EPSCs, vesicle pool parameters, and presynaptic $Ca^{2+}$ currents at hippocampal mossy fiber–CA3 pyramidal neuron synapses.

**Table S2.**

| **Parameter [unit]** | **Mean ± SEM / median ($25^{th}$–$75^{th}$ percentile)** | **p-value and statistical test** | **Fig.** |
|---|---|---|---|
| mEPSC frequency [Hz] | Ctrl: 1.84 ± 0.31 / 1.29 (0.80–2.62) | $p < 0.001$; Wilcoxon signed rank test | 1L |
| | Bac: 0.63 ± 0.09 / 0.54 (0.27–0.89) | | |
| mEPSC peak amplitude median [pA] | Ctrl: 31.34 ± 1.50 / 29.18 (25.82–36.94) | $p = 0.004$; Wilcoxon signed rank test | 1M |
| | Bac: 27.75 ± 1.26 / 27.42 (23.87–32.84) | | |
| mEPSC frequency [Hz] (rise time < 1 ms) | Ctrl: 1.48 ± 0.29 / 1.00 (0.59–2.05) | $p < 0.001$; Wilcoxon signed rank test | S2C |
| | Bac: 0.48 ± 0.07 / 0.44 (0.20–0.60) | | |
| mEPSC peak amplitude median [pA] (rise time < 1 ms) | Ctrl: 34.32 ± 1.28 / 32.16 (30.43–39.15) | $p = 0.015$; Wilcoxon signed rank test | S2D |
| | Bac: 31.21 ± 1.05 / 30.37 (28.17–35.84) | | |

Effects of 20 µM baclofen on miniature EPSCs in CA3 pyramidal neurons.

**Table S3.**

| **Parameter [unit]** | **Mean ± SEM / median ($25^{th}$–$75^{th}$ percentile)** | **p-value and statistical test** | **Fig.** |
|---|---|---|---|
| $EPSP_1$ peak amplitude [mV] | Ctrl: 8.75 ± 1.79 / 10.55 (3.40–12.31) | $p$ = 0.008; Wilcoxon signed rank test | 2B |
| | Bac: 0.54 ± 0.17 / 0.36 (0.14–1.05) | | |
| $EPSP_1$ failure proportion | Ctrl: 8.75 ± 4.41 / 0 (0–20) | $p$ = 0.008; Wilcoxon signed rank test | 2C |
| | Bac: 55 ± 9.06 / 55 (40–77.50) | | |
| Probability of $1^{st}$ AP | Ctrl: 0.16 ± 0.11 / 0 (0–0.25) | $p$ = 0.454; ART ANOVA with contrast test (Holm adjustment) | 2D |
| | Bac: 0 ± 0 / 0 (0–0) | | |
| Probability of $2^{nd}$ AP | Ctrl: 0.49 ± 0.15 / 0.50 (0.03–0.95) | $p$ = 0.001; ART ANOVA with contrast test (Holm adjustment) | |
| | Bac: 0 ± 0 / 0 (0–0) | | |
| Probability of $3^{rd}$ AP | Ctrl: 0.66 ± 0.15 / 0.90 (0.18–0.90) | $p$ = 0.006; ART ANOVA with contrast test (Holm adjustment) | |
| | Bac: 0.08 ± 0.05 / 0 (0–0.15) | | |

Effects of 20 µM baclofen on evoked EPSPs and action potential firing at hippocampal mossy fiber–CA3 pyramidal neuron synapses.

**Table S4.**

| Parameter [unit] | Mean ± SEM / median (25$^{th}$–75$^{th}$ percentile) | P-value and statistical test | Fig. |
|---|---|---|---|
| % PTP | HFS: 458.89 ± 166.75/ 219.61 (165.72–638.73) | HFS vs. HFS+LS (5 Hz): $p = 0.008$; HFS vs. HFS+LS (20 Hz): $p = 0.036$; Kruskal-Wallis test with Dunn's multiple comparisons test | 9G |
| | HFS+LS (5 Hz):120.59 ± 16.37 / 131.45 (84.45–159.64) | | |
| | HFS+LS (20 Hz):142.65 ± 23.67 / 140.88 (108.12–175.60) | | |
| $EPSC_1$ peak amplitude [pA] | Ctrl: 193.04 ± 71.95 / 107.14 (37.61–289.11) | $p = 0.002$; Wilcoxon signed rank test | 9H |
| | HFS: 433.28 ± 98.54 / 393.34 (181.16–630.83) | | |
| | Ctrl: 247.26 ± 50.37 / 242.00 (137.00–307.21) | $p = 0.109$; Wilcoxon signed rank test | |
| | HFS+LS (5 Hz): 332.33 ± 88.97 / 347.36 (78.47–458.08) | | |
| | Ctrl: 251.63 ± 89.95 / 127.00 (81.92–334.51) | $p = 0.105$; Wilcoxon signed rank test | |
| | HFS+LS (20 Hz): 307.03 ± 100.05 / 235.61 (80.78–433.63) | | |
| $EPSC_2$ / $EPSC_1$ | Ctrl: 2.66 ± 0.55 / 1.85 (1.40–4.00) | $p = 0.006$; ART ANOVA with contrast test (Holm adjustment) | S8A |
| | HFS: 1.33 ± 0.39 / 0.98 (0.76–1.18) | | |
| $EPSC_3$ / $EPSC_1$ | Ctrl: 3.49 ± 1.02 / 2.33 (1.62–4.54) | $p = 0.002$; | |

| | | | |
|---|---|---|---|
| | HFS: 1.32 ± 0.32 / 0.93 (0.73–1.81) | ART ANOVA with contrast test (Holm adjustment) | |
| $EPSC_2$ / $EPSC_1$ | Ctrl: 1.19 ± 0.17 / 1.19 (0.79–1.42) | $p = 0.483$; ART ANOVA with contrast test (Holm adjustment) | |
| | HFS+LS (5 Hz): 0.83 ± 0.21 / 0.73 (0.27–1.26) | | |
| $EPSC_3$ / $EPSC_1$ | Ctrl: 1.40 ± 0.23 / 1.65 (0.63–1.95) | $p = 0.997$; ART ANOVA with contrast test (Holm adjustment) | |
| | HFS+LS (5 Hz): 1.33 ± 0.34 / 0.94 (0.61–2.38) | | |
| $EPSC_2$ / $EPSC_1$ | Ctrl: 1.78 ± 0.30 / 1.32 (1.13–2.60) | $p = 0.572$; ART ANOVA with contrast test (Holm adjustment) | |
| | HFS+LS (20 Hz): 1.32 ± 0.30/ 1.12 (0.63–1.86) | | |
| $EPSC_3$ / $EPSC_1$ | Ctrl: 2.42 ± 0.81 / 1.63 (1.27–2.46) | $p = 0.572$; ART ANOVA with contrast test (Holm adjustment) | |
| | HFS+LS (20 Hz): 1.47 ± 0.34 / 1.30 (0.84–2.18) | | |
| RRP [nA] | Ctrl: 1.24 ± 0.27 / 1.07 (0.68–1.53) | $p = 0.002$; Wilcoxon signed rank test | 10D |
| | HFS: 2.02 ± 0.32 / 1.83 (1.24–2.44) | | |
| | Ctrl: 1.17 ± 0.19 / 1.20 (0.78–1.68) | $p = 0.641$; Wilcoxon signed rank test | |
| | HFS+LS (5 Hz): 1.23 ± 0.30 / 1.29 (0.42–1.85) | | |
| | Ctrl: 1.26 ± 0.32 / 0.96 (0.55–1.57) | $p = 0.160$; Wilcoxon signed rank test | |

| | | | |
|---|---|---|---|
| | HFS+LS (20 Hz): 1.46 ± 0.27 /<br>1.14 (0.92–2.05) | | |
| Refilling rate [pA ms$^{-1}$] | Ctrl: 4.63 ± 0.76 /<br>3.74 (3.18–6.50) | *p* = 0.002;<br>Wilcoxon signed rank test | 10E |
| | HFS: 2.72 ± 0.65 /<br>1.81 (1.10–4.48) | | |
| | Ctrl: 3.54 ± 0.83 /<br>3.30 (2.26–4.12) | *p* = 0.461;<br>Wilcoxon signed rank test | |
| | HFS+LS (5 Hz): 3.36 ± 0.83 /<br>3.38 (1.76–3.45) | | |
| | Ctrl: 5.81 ± 1.40 /<br>5.26 (2.17–8.44) | *p* = 0.309;<br>Wilcoxon signed rank test | |
| | HFS+LS (20 Hz): 6.31 ± 1.59 /<br>5.84 (2.28–8.91) | | |
| $P_v$ | Ctrl: 0.13 ± 0.03 /<br>0.11 (0.09–0.16) | *p* = 0.014;<br>Wilcoxon signed rank test | 10F |
| | HFS: 0.20 ± 0.02 /<br>0.22 (0.13–0.25) | | |
| | Ctrl: 0.22 ± 0.04 /<br>0.17 (0.15–0.34) | *p* = 0.039;<br>Wilcoxon signed rank test | |
| | HFS+LS (5 Hz): 0.29 ± 0.05 /<br>0.25 (0.17–0.43) | | |
| | Ctrl: 0.17 ± 0.02 /<br>0.18 (0.12–0.21) | *p* = 0.922;<br>Wilcoxon signed rank test | |
| | HFS+LS (20 Hz): 0.17 ± 0.03 /<br>0.17 (0.08–0.25) | | |
| RRP (TR method) [nA] | Ctrl: 1.50 ± 0.28 /<br>1.36 (0.96–1.74) | *p* = 0.004;<br>Wilcoxon signed rank test | S8C |
| | HFS: 2.24 ± 0.32 / | | |

| | | | |
|---|---|---|---|
| | 2.20 (1.42–2.59) | | |
| $P_v$ (TR method) | Ctrl: 0.12 ± 0.02 / 0.12 (0.07–0.15) | $p$ = 0.027; Wilcoxon signed rank test | S8D |
| | HFS: 0.19 ± 0.02 / 0.19 (0.13–0.24) | | |

Effects of synaptically released GABA on post-tetanic potentiation and vesicle pool parameters at hippocampal mossy fiber–CA3 pyramidal neuron synapses.

**Table S5.**

| Biophysical effects | Mean ± SEM | Fig. |
|---|---|---|
| Block of EPSC | 89.9 ± 2.1% | 1D |
| Reduction of RRP | 76.0 ± 6.2% | 3B |
| Reduction of $P_v$ | 48.4 ± 8.0% | 3C |
| Predicted reduction of release<br>$1 - (RRP_{Bac} / RRP_{Ctrll} \times P_{v, Bac} / P_{r, Ctrl})$ | 87.1% | N/A |
| Reduction of $Ca^{2+}$ current | 35.0 ± 3.8% | 3G |
| Predicted reduction of $P_v$<br>$1 - ((I_{Ca, Bac} / I_{Ca, Ctrl})^m$ * | 52.8% | N/A |

Summary of effects of 20 μM baclofen and quantitative prediction. *Using experimentally determined power coefficient of release (1.77; (*52*)).

**Table S6.**

| **Parameter [unit]** | **Median ($25^{th}$–$75^{th}$ percentile) / mean ± SEM** | **p-value and statistical test** | **Fig.** |
|---|---|---|---|
| 2D EM number for docked clear vesicles (0–2 nm) per 100 nm AZ profile length | Ctrl: 0.43 (0.31–0.58) | *p* < 0.001; Mann–Whitney U test | 4C |
| | Bac: 0.25 (0–0.45) | | |
| 2D EM diameter of docked** clear vesicles [nm] | Ctrl: 48.10 (40.90–67.81) | *p* = 0.040; Mann–Whitney U test | 4E |
| | Bac: 43.46 (40.75–54.52) | | |
| 2D EM diameter of docked** clear vesicles cut-off < 65 nm [nm] | Ctrl: 43.88 (38.84–48.38) | *p* = 0.273; Mann–Whitney U test | 4E |
| | Bac: 41.99 (38.70–46.41) | | |
| 3D ET number for total vesicle pool (0–50 nm) per 0.01 µm2 AZ area | Ctrl: 3.07 (2.27–4.11) | *p* = 0.248; Mann–Whitney U test | 5E |
| | Bac: 2.74 (1.92–3.36) | | |
| 3D ET number for docked clear vesicles (0–2 nm) per 0.01 $\mu m^2$ AZ area | Ctrl: 1.10 (0.92–1.54) | *p* = 0.024; Mann–Whitney U test | 5F, S6C |
| | Bac: 0.70 (0.49–1.22) | | |
| Mean NND docked clear vesicles [nm] | Ctrl: 97.38 (68.96–143.98) | *p* = 0.023; Mann–Whitney U test | 5H |
| | Bac: 124.77 (72.43–173.67) | | |
| Distance of docked clear vesicles (0–2 nm) to AZ COM [nm] | Ctrl: 93.88 (50.40–134.85) | *p* = 0.681; Mann–Whitney U test | 5K |
| | Bac: 85.97 (49.82–135.47) | | |
| AZ surface area [$\mu m^2$] | Ctrl: 0.042 (0.031–0.065) | *p* = 0.058; Mann–Whitney U test | S6B |
| | Bac: 0.059 (0.041–0.083) | | |

| | | | |
|---|---|---|---|
| 3D ET number for docked clear vesicles (0–5 nm) per 0.01 μm$^2$ | Ctrl: 1.31 (1.01–1.89)<br>Bac: 1.03 (0.77–1.53) | $p = 0.101$; Mann–Whitney U test | S6C |
| 3D ET number for docked clear vesicles (2–5 nm) per 0.01 μm$^2$ AZ area | Ctrl: 0.23 (0–0.31)<br>Bac: 0.22 (0.07–0.41) | $p = 0.662$; Mann–Whitney U test | S6C |
| 3D ET docked (0–2 nm) clear vesicle diameter [nm] | Ctrl: 49.63 (45.71–73.73)<br>Bac: 51.03 (45.34–65.75) | $p = 0.917$; Mann–Whitney U test | S6D |
| 3D ET docked (0–5 nm) clear vesicle diameter [nm] | Ctrl: 49.47 (45.87–66.38)<br>Bac: 48.21 (43.99–54.56) | $p = 0.113$; Mann–Whitney U test | S6D |
| 3D ET docked (2–5 nm) clear vesicle diameter [nm] | Ctrl: 49.04 ± 1.09<br>Bac: 45.06 ± 0.94 | $p = 0.009$; Student's t-test | S6D |
| Total vesicle volume per 0.01 μm$^2$ AZ area [nm$^3$] | Ctrl: 126264.05 (110861.36–224079.49)<br>Bac: 81091.42 (34553.49–155805.94) | $p = 0.004$; Mann–Whitney U test | S6E |
| 3D ET number of docked dense-core vesicles (0–2 nm) per 0.01 μm$^2$ AZ area | Ctrl: 0.167 (0–0.310)<br>Bac: 0.200 (0–0.341) | $p = 0.972$; Mann–Whitney U test | S6G |
| 3D ET number of total dense-core vesicles (0–50 nm) per 0.01 μm$^2$ AZ area | Ctrl: 0.29 ± 0.06<br>Bac: 0.29 ± 0.05 | $p = 0.941$; Student's t-test | S6G |
| | Ctrl: 76.71 ± 2.13 | $p = 0.003$; | S6H |

| | | | |
|---|---|---|---|
| 3D ET dense-core vesicle diameter [nm] | Bac: 68.32 ± 1.66 | Student's t-test | |
| Total dense core vesicle volume per 0.01 μm$^2$ AZ area [nm$^3$] | Ctrl: 74417.94 ± 15125.92 | *p* = 0.424; Student's t-test | S6I |
| | Bac: 59737.36 ± 8805.55 | | |
| Mean NND all vesicles (0–50 nm) [nm] | Ctrl: 53.50 (45.07–65.59) | *p* = 0.395; Mann–Whitney U test | N/A |
| | Bac: 54.73 (43.76–73.56) | | |

Effects of 20 μM baclofen on ultrastructure of hippocampal mossy fiber–CA3 synapses.
**Docked vesicle analysis was conducted applying the definition of vesicle docking (0–2 nm) according to (*35*, *53*), except for the 2D TEM diameter quantification (0–5 nm) (*26*).

**Table S7.**

| **Parameter [unit]** | **Median ($25^{th}$–$75^{th}$ percentile)** | **p-value and test** | **Fig.** |
|---|---|---|---|
| ***LS (5 Hz)*** | | | |
| 2D EM number for docked clear vesicles (0–2 nm) per 100 nm AZ profile length | non-LS: 0.66 (0.51–0.81) | $p < 0.001$; Mann–Whitney U test | 8D |
| | LS (5 Hz): 0.31 (0–0.49) | | |
| ***LS (20 Hz)*** | | | |
| 2D EM number for docked clear vesicles (0–2 nm) per 100 nm AZ profile length | non-LS: 0.60 (0.49–0.84) | $p < 0.001$; Mann–Whitney U test | 8D |
| | LS (5 Hz): 0.30 (0.15–0.48) | | |
| ***LS (20 Hz) + gabazine*** | | | |
| 2D EM number for docked clear vesicles (0–2 nm) per 100 nm AZ profile length | non-LS: 0.68 (0.53–0.84) | $p < 0.001$; Mann–Whitney U test | 8D |
| | LS (20 Hz + gabazine): 0.34 (0.21–0.43) | | |

Effects of synaptically released GABA on ultrastructure of hippocampal mossy fiber–CA3 synapses.
